# Regulation of the desmosome-intermediate filament linkage enables an adaptive mechano-response within the stratified epidermis

**DOI:** 10.64898/2026.08.28.747586

**Authors:** Abbey L Perl, Rosemary DiDominicis, Joshua Broussard, Constadina Arvanitis, Kathleen J Green

**Affiliations:** Department of Pathology, Feinberg School of Medicine, Northwestern University, Chicago, Illinois; Department of Dermatology, Feinberg School of Medicine, Northwestern University, Chicago, Illinois; Center for Advanced Microscopy, Feinberg School of Medicine, Northwestern University, Chicago, Illinois; Robert H. Lurie Comprehensive Cancer Center, Northwestern University, Chicago, Illinois

## Abstract

Skin, the body’s largest mechanosensitive organ, relies on a tension gradient across epidermal layers to maintain structure and function, but how mechanical force contributes to epidermal development and disease pathogenesis is poorly understood. By anchoring intermediate filaments (IF) to the plasma membrane, desmosomes, the most abundant intercellular junctions in the epidermis, help create a supracellular scaffolding that provides mechanical resilience to the tissue. However, the contribution of the desmosome-IF network to the epidermal response to mechanical strain remains unknown. Here we show that the desmosome-IF connection is not only required to induce a proper cellular mechano-response but is actively strengthened in response to stretch through the PP2A-mediated phospho-regulation of the cytoskeletal linker protein desmoplakin (DP). Additionally, we show in human skin dephosphorylated DP localizes to high tension layers, suggesting this mechano-response mechanism is coordinated with the epidermal tension gradient. Furthermore, in models of Carvajal syndrome, a cardio-cutaneous disorder caused by truncating DP mutations, cells lose mechano-responsive behavior and exhibit abnormal morphology in high-tension epidermal layers. Together, these findings identify the DP-IF network as a key component of the response to mechanical strain and show that its disruption compromises epidermal homeostasis and contributes to disease pathogenesis.

## INTRODUCTION

The skin is a highly mechanosensitive organ as it experiences mechanical changes on both a macro and micro scale. It is forced to respond and adapt to large-scale mechanical changes that occur during development and as a result of physical activity (Hsu et al., 2018). Additionally, the skin experiences changes in its micro-mechanical environment including that conferred by an intrinsic tension gradient that exists across the multi-layered epidermis, forcing cells to mechanically adapt as they move upwards through the tissue (Broussard et al., 2021; Fiore et al., 2020; Mao and Wickstrom, 2024). The epidermal mechanical gradient spans the basal-apical axis where the mechanics change from compressive forces in the basal layer to high-tension in the apical layers corresponding with the tight junction formation and increased cortical actin. The skin responds to its mechanical environment through altered signaling (Mao and Wickstrom, 2024; Rubsam et al., 2023; Villeneuve et al., 2024). This includes a well-defined proliferative response to prolonged stretching that facilitates tissue expansion, as well as more immediate responses within individual keratinocytes (Biggs et al., 2020).

Epidermal keratinocytes’ response to external mechanical strain begins with an immediate cytoskeletal response that leads to a subsequent reorganization of cell morphology. Similar to other epithelia, within hours of experiencing mechanical stretch keratinocytes align their actin and intermediate filament (IF) cytoskeletal networks and elongate their cell shapes (De et al., 2008; Fudge et al., 2008; Lien and Wang, 2021; Noethel et al., 2018). Functional studies revealed that stretch-induced actin reorientation is calcium dependent and requires intact actin-associated junctions, including cell-substrate focal adhesions and cell-cell adherens junctions (Noethel et al., 2018). These cytoskeletal and cellular reorientation responses help mitigate the intracellular stress induced by mechanical strain (Le et al., 2016; Nava et al., 2020).

Though these findings support an important role for calcium-dependent adhesions in facilitating the cells’ adaptative response to mechanical strain, how desmosomes, the most prominent calcium-dependent adhesive structures in the epidermis, contribute to strain adaptation is poorly understood. Similar to adherens junctions in their molecular blueprint, desmosomes are built from two desmosomal cadherin subclasses, desmogleins and desmocollins, whose cytoplasmic tails recruit intracellular plaque proteins that link IFs to the plasma membrane (Perl et al., 2024). The desmosome-IF network facilitates the formation of a supracellular scaffolding that is critical for maintaining the mechanical resilience of the tissue (Broussard et al., 2020; Fudge et al., 2008; Hatzfeld et al., 2017; Jin et al., 2021). Highlighting the crucial role desmosomes have in maintaining tissue adhesions, disruption of desmosomes is associated with diseases in tissues that undergo high levels of mechanical strain, including the skin and heart (Mezzano and Sheikh, 2012; Perl et al., 2024; Schmidt and Koch, 2007).

While regulation of intracellular mechanics has been attributed largely to adherens junctions and the associated actin cytoskeleton, a role for the desmosome-IF network in modulating a cells’ mechanical properties is beginning to emerge (Broussard et al., 2021). In particular, the desmosome-IF linker protein desmoplakin (DP) can modulate the strength of the DP-IF connection to impact actin-dependent forces, resulting in altered cell adhesion and stiffness (Broussard et al., 2017). The DP-IF connection is regulated through GSK3-dependent phosphorylation of a 20-amino acid Ser-Gly-Arg phospho-motif at DP’s C-terminus that impacts DP’s affinity for IF (Albrecht et al., 2015; Broussard et al., 2017; Stappenbeck et al., 1994). Phosphorylation of this motif weakens the DP-IF linkage. Conversely, we showed that PP2A-B55α mediated dephosphorylation of DP strengthens the DP-IF connection and bolsters keratinocytes’ adhesive capacity (Godsel et al., 2005; Hobbs and Green, 2012; Perl et al., 2023). Moreover, expression of constitutively hypo-phosphorylated DP C-terminus results in stiffer cells with increased intercellular forces and has been shown to be protective in skin disease models (Broussard et al., 2017; Dehner et al., 2014).

Given the importance of the DP-IF interaction in tuning actin-driven cell mechanics, we set out to investigate the role of the desmosome-IF connection in facilitating keratinocytes’ mechano-response within the stratified epidermis. We show that the DP-IF connection is not only required for a proper keratinocyte mechanical response but is actively regulated through phosphatase-mediated DP dephosphorylation in response to stretch. Additionally, we demonstrate that the phosphatase-dependent mechano-response is required to maintain the cells’ adhesive capacity in response to mechanical strain. In human skin samples we identify an inverse correlation between DP’s phospho-regulation and the epidermal mechanical gradient. Moreover, DP mutations responsible for human Carvajal Syndrome lead to a disrupted mechano-response to stretch, resulting in defects in cell shape polarization in the high-tension layers of Carvajal mice. Based on these findings, we propose a model whereby keratinocytes actively regulate the DP-IF connection through DP dephosphorylation in response to changes in the mechanical environment to maintain the skin’s homeostatic tissue architecture.

## RESULTS

### The desmoplakin-IF connection is required for keratinocytes’ cellular response to stretch

To determine how the desmosome-IF connection contributes to keratinocytes’ response to mechanical strain, we developed DP Crispr knockout cells in N/TERT-2G immortalized keratinocytes (Dickson et al., 2000). Utilizing guide RNAs targeting a negative control sequence or the start codon of the desmoplakin gene (*DSP*), we generated two stable lines from each condition: crControl-1, crControl-2, cr*DSP*-1, cr*DSP*-2. DP knockout was confirmed by sequencing and protein expression (Supplemental Figure 1).

Using a Flex Cell vacuum-based stretcher, cells were subjected to 18% cyclic strain for 4 or 24 hours. After both 4 and 24 hours, crControl cells exhibited robust cell shape changes compared to unstretched controls (Figure 1A, Supplemental Figure 2A). Cell shapes were quantified using aspect ratio (AR), feret, and roundness measurements, which consistently showed crControl cells become elongated and less round with stretch (Figure 1B-C, Supplemental Figure 2C). Additionally, cell angle measurements showed that cell elongation occurs perpendicular to the direction of strain (Supplemental Figure 2B). Conversely, DP knockout cells subjected to stretch exhibited no significant changes in their cell morphology (Figure 1A-C, Supplemental Figure 2A-C). These data suggest that DP is necessary for the cell’s mechanical response to stretch.

**Figure 1:**
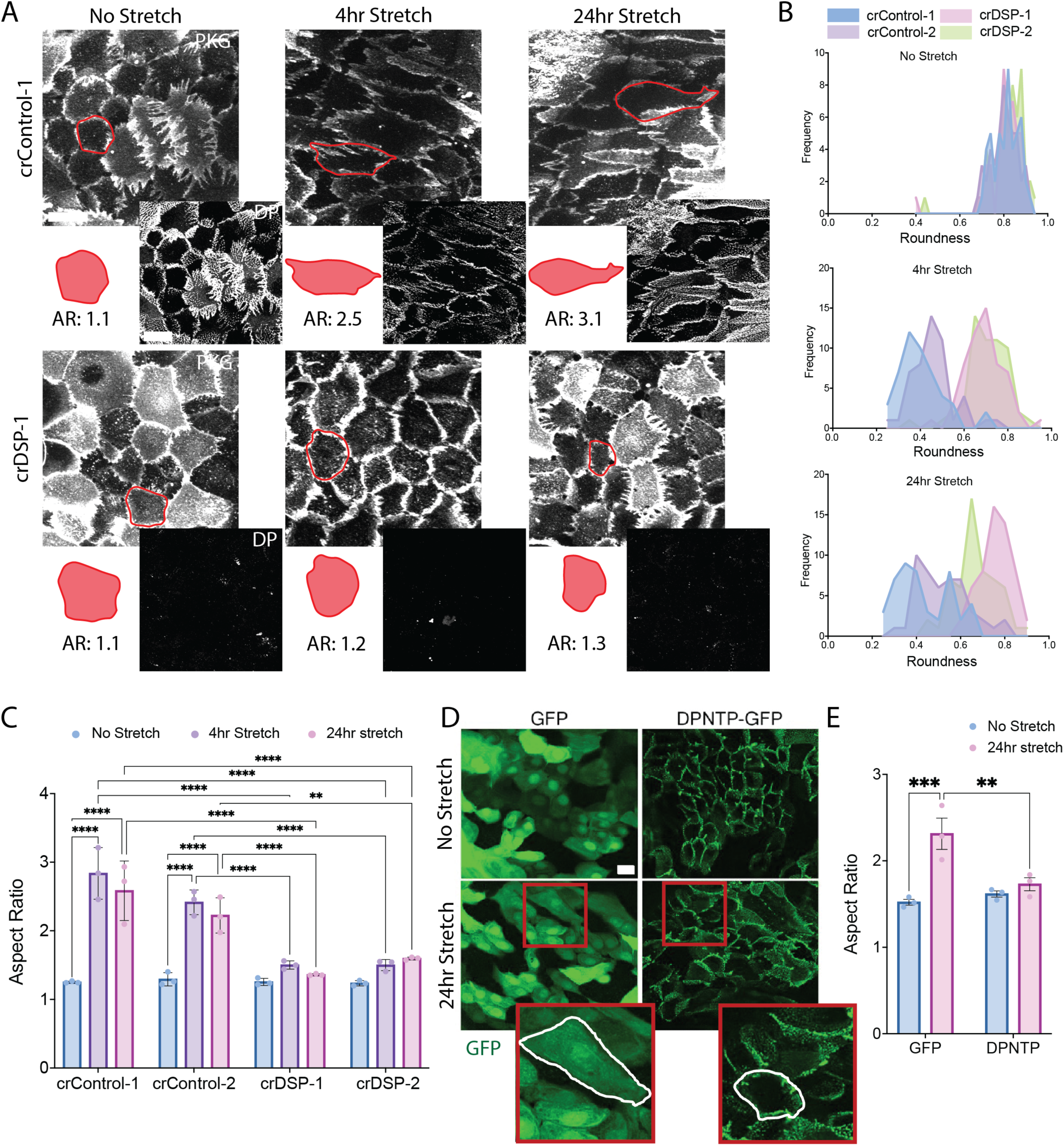
The desmosome-cytoskeletal linkage is required for stretch-induced shape changes. A) DP knockout cells were stretched for 4 and 24 hours. Immunofluorescence images of crControl-1 (top) and cr*DSP*-1 (bottom) cells post treatment detecting PKG and DP. Representative cells are outlined in red and reconstructed below image. AR values of the representative cell are depicted below. B) Frequency plots representing roundness measurements of individual cells from each condition in (A). Plots are separated by treatment. C) Mean AR values from 3 biological replicates of (A). D) NHEKs transduced with GFP or DPNTP were stretched for 24 hours. Immunofluorescence images from cells after stretching detecting GFP expression. E) Mean aspect ratio values from 3 biological replicates of (D). Images were acquired with an apotome confocal. Statistical analyses are from a two-way Anova with multiple comparisons from an n=3. Scale bars are 20 μm. P-values are represented by (*) = 0.05, (**) = 0.01, (***) = 0.001, (****) = 0.0001.

Given the essential role of DP as the cytoskeletal linker between the desmosome and IF network, we next addressed whether the DP-IF connection was facilitating the DP-dependent mechanical stretch response. Toward this end, we used a dominant negative N-terminal truncated form of DP (DPNTP-GFP) lacking the IF binding domain expressed in primary human epidermal keratinocytes (NHEKs) to test whether loss of the DP-IF connection interfered with the cells’ mechano-response (Bornslaeger et al., 1996; Huen et al., 2002). Transduced keratinocytes were stretched for 24 hours resulting in elongation of GFP control cells. However, no significant cell shape response was seen in DPNTP expressing cells demonstrating that the DP-IF connection is important in the cellular response to stretch (Figure 1D-E, Supplemental Figure 2D).

### Stretch-induced actin reorientation is unaffected by the DP-IF linkage

Previous reports showed that actin dynamics and actin-associated junctions contributed to the stretch-induced reorientation response (Noethel et al., 2018). Therefore, to understand how the actin cytoskeleton is affected by loss of the DP-IF connection, stretched GFP and DPNTP-GFP expressing keratinocytes were stained with phalloidin to visualize actin filaments. In both GFP and DPNTP conditions, stretch induced alignment of the actin filaments as demonstrated by actin fiber anisotropy (Figure 2A-B). Similarly, the actin filaments of crControl and cr*DSP* cells stretched for 4 hours were significantly aligned despite the lack of cell shape changes in the cr*DSP* cells (Figure 2C-D). These data suggest that in the absence of an intact DP-IF connection, actin re-alignment is not sufficient to induce the cell shape response to mechanical stretch.

**Figure 2:**
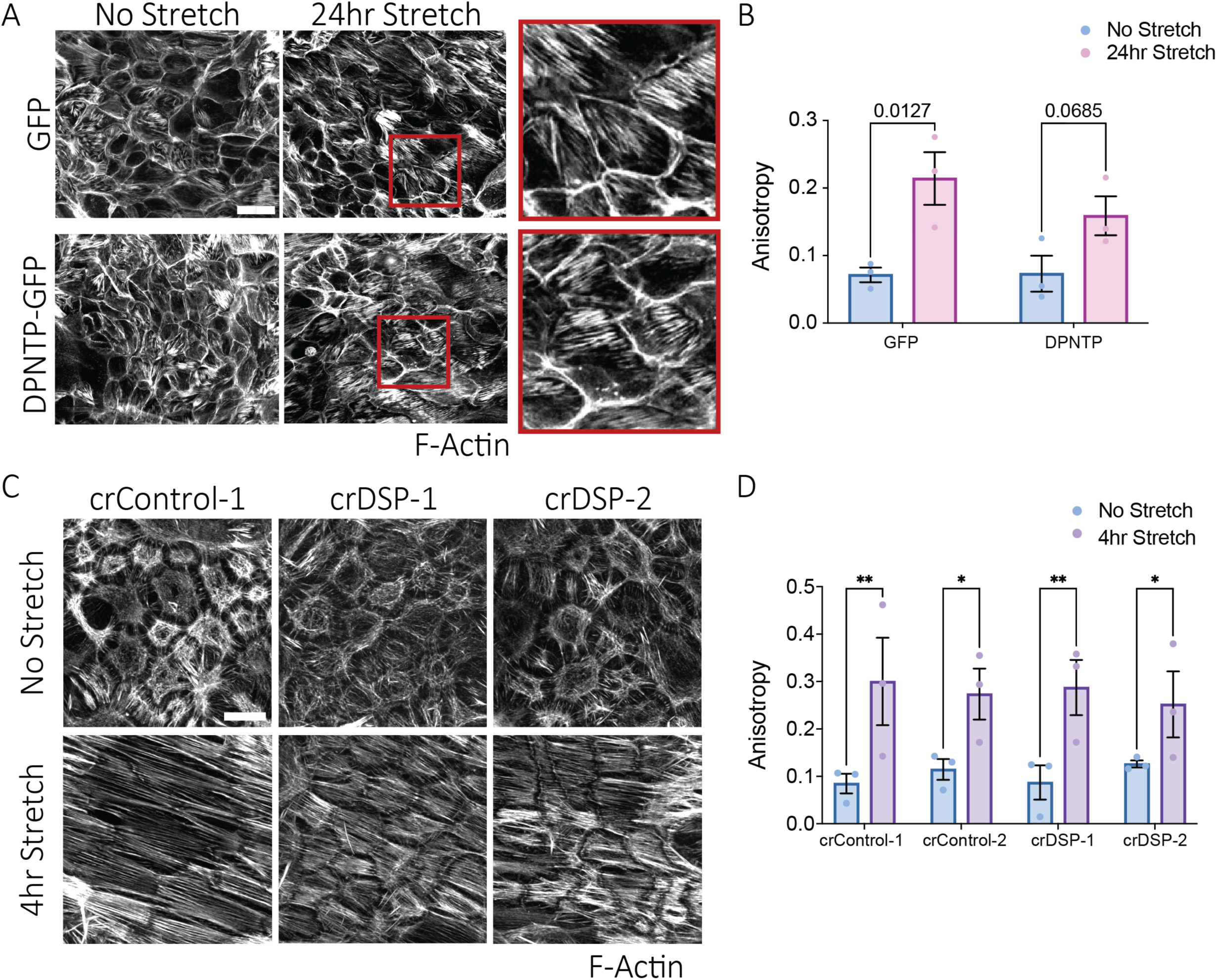
Stretch-induced actin reorientation is unaffected by the DP-IF linkage. *A) NHEKs* transduced with GFP or DPNTP were stretched for 24 hours. Images show immunofluorescence staining with phalloidin from cells after stretching detecting actin filaments. Insets are taken from 24 hour stretch conditions. B) Graph of the mean actin fiber anisotropy values from 3 biological replicates from (A). C) DP knockout cells were stretched for 4 hours. Images show immunofluorescence staining with phalloidin from cells after stretching detecting actin filaments. D) Graph of the mean actin fiber anisotropy values from 3 biological replicates from (C). Scale bars are 20 μm. Images were acquired with a spinning disk confocal. Statistical analyses are from a two-way Anova with multiple comparisons from an n=3. P-values are represented by (*) = 0.05, (**) = 0.01, (***) = 0.001, (****) = 0.0001.

### Stretch-induced shape changes require DP phospho-regulation

Our previous work identified a phospho-regulatory motif on the C-terminus of DP that tunes DP-IF binding affinity (Stappenbeck et al., 1994). Specifically, hyper-phosphorylation of this motif results in weakened DP-IF binding (Albrecht et al., 2015). Conversely, hypo-phosphorylated DP results in ultra strong DP-IF binding and increased cell adhesion strength (Godsel et al., 2005; Hobbs and Green, 2012; Stappenbeck et al., 1994). To determine whether DP-IF connections are regulated through modification of this motif during stretch, NHEKs were stretched for 4 hours and stained for total DP alongside phospho-S2849-DP (pDP), the priming phospho-site for the C-terminus phospho-regulatory motif (Figure 3A)(Albrecht et al., 2015). Staining showed a significant loss of pDP at cell membranes after 4 hours of stretch, suggesting a potential strengthening of the DP-IF connections in response to mechanical strain (Figure 3B-C). In conjunction with decreased pDP, keratin IF became aligned along the long axis of cells at 4 hours of stretch (Figure 3D).

**Figure 3:**
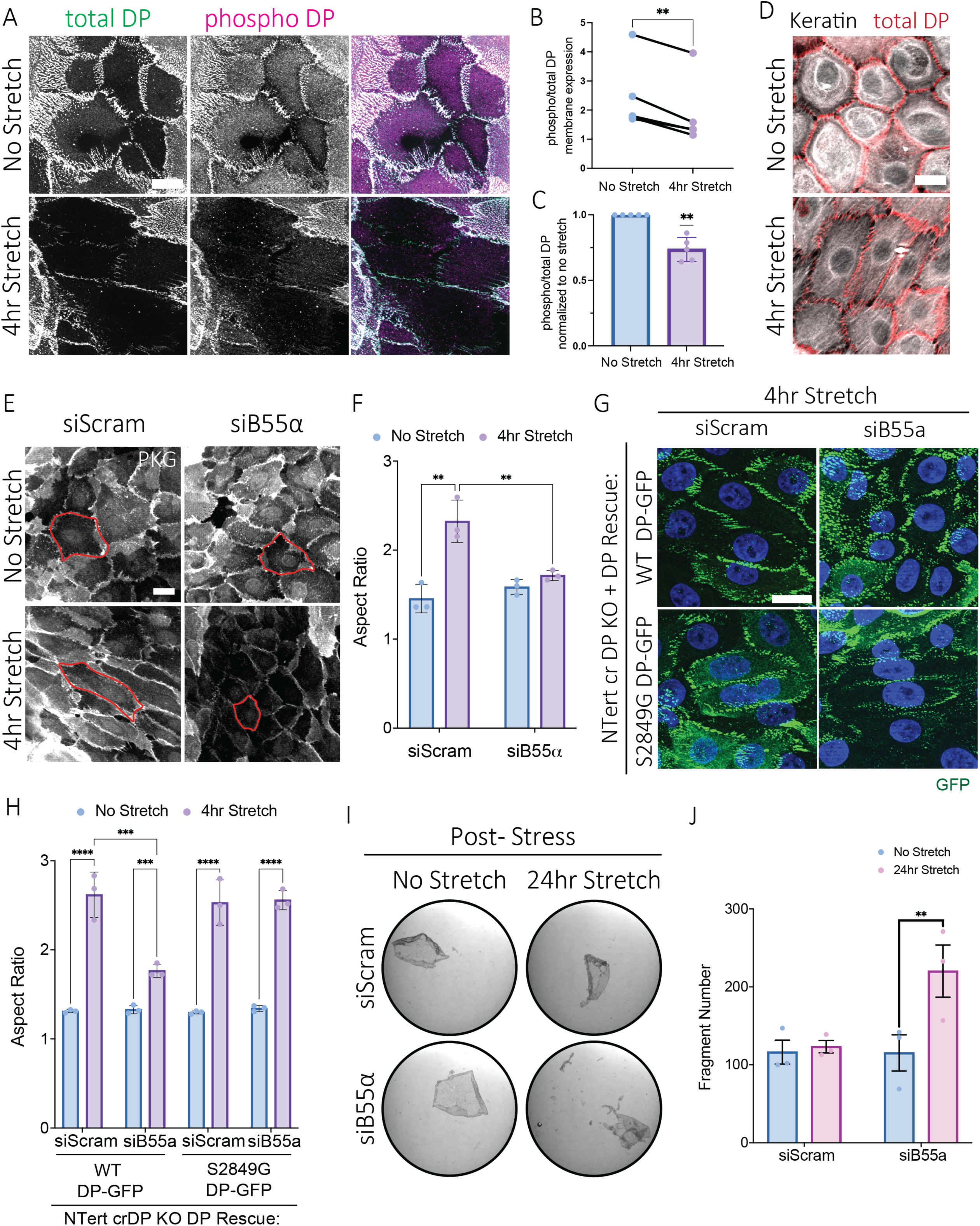
Stretch-induced PP2A-B55α mediated dephosphorylation of DP facilitates keratinocytes mechano-response and reinforced epidermal adhesive capacity. A) Images show immunofluorescence staining from NHEKs stretched for 4 hours. Images were acquired with an apotome confocal. Representative images from no stretch and 4hr stretched conditions staining for total DP (left, green) and phospho-S2849-DP (middle, magenta) with the overlayed staining (right). B-C) Staining intensity at cell membranes were measured using line scans of (A). Graphs show phospho-DP: total DP ratios of mean values from 5 biological replicates depicting the raw values (C) or the values normalized to each no stretch control (B). D) Images show immunofluorescence staining acquired on a spinning disk confocal from NHEKs stretched for 4 hours and stained for keratin 14 (white) or total DP (red). E) NHEK cells with B55α knockdown via targeted siRNA were stretched for 4 hours. Representative images from no stretch and 4hr stretched conditions stained for PKG are shown to depict cell shapes. Images were acquired with an apotome confocal. Representative cells are outlined in red. F) Mean aspect ratio values from 3 biological replicates from (E). G) DP knockout cells cr*DSP*-1 were transduced with either WT-DP-GFP or S2849G-DP-GFP and B55α was knockdown in both conditions via targeted siRNA. Images were acquired with an apotome confocal. Representative images of cells after 4 hours of stretch and stained for GFP are shown. H) Graph of mean aspect ratio values from 3 biological replicates from (G). I) NHEK cells with B55α knockdown via targeted siRNA were stretched for 24 hours and treated with dispase to lift cell sheets and subjected to mechanical stress. Images of cell sheets after a dispase assay mechanical stress. J) Cell sheet fragments from (I) were quantified from 3 biological replicates. Scale bars are 20 μm. Statistical analyses are from a two-way Anova with multiple comparisons from an n=3-5. P-values are represented by (*) = 0.05, (**) = 0.01, (***) = 0.001, (****) = 0.0001.

To understand the role of hypo-phosphorylated DP in the mechano-response, DP rescue experiments were performed in the cr*DSP*-1 cells. cr*DSP*-1 cells were transduced with either a GFP control, wild-type-DP-GFP (WT-DP-GFP), or a mutant S2849G-DP-GFP which was previously shown to act as a constitutively hypo-phosphorylated DP (Supplemental Figure 3). Transduced cells were stretched for 4 hours alongside crScram-1 cells. GFP staining and immunoblotting confirmed rescue and proper localization of WT-DP-GFP and S2849G-DP-GFP protein in transduced cells (Supplemental Figure 3A-B). Cell shape changes were quantified using plakoglobin (PKG) as a cell membrane marker (Supplemental Figure 3C). Expression of the S2849G-DP-GFP was sufficient to restore the stretch-induced cell shape changes as efficiently as WT-DP-GFP or the crScram-1 cells as measured by AR, feret or roundness (Supplemental Figure 3D-G). These data suggest that hypo-phosphorylated DP is sufficient for the cellular response to mechanical stretch.

We previously identified the Protein Phosphatase 2A (PP2A) with the B55α regulatory subunit as the phosphatase responsible for dephosphorylating DP’s C-terminus (Perl et al., 2023). Therefore, we sought to determine if PP2A-B55α regulation of DP is important in the cellular response to stretch. To address this, we depleted the PP2A regulatory subunit gene B55α in NHEKs with siRNA and subjected cells to 4 hours of cyclic stretch (Figure 3E). B55α knockdown impaired the cell elongation response seen in scramble control cells as determined by AR, feret, and roundness measurements (Figure 3F, Supplemental Figure 4A-D). These data suggest that PP2A-B55α contributes to the cells’ response to stretch.

To determine whether PP2A-B55α’s regulation of the cells’ mechano-response is through phospho-regulation of DP’s C-terminus, cr*DSP*-1 cells were transduced with either WT-DP-GFP or the constitutively hypo-phosphorylated S2849G-DP-GFP. Both WT-and S2849G-DP-GFP expressing cells were treated with B55α targeting siRNA and subjected to 4 hours of stretch alongside scramble and unstretched controls (Figure 3G, Supplemental Figure 4E). Under these conditions, B55α loss significantly impaired the cells response to stretch in the presence of WT-DP-GFP (Figure 3H, Supplemental Figure 4E-G). These data confirm that WT-DP-GFP rescue in the cr*DSP* cells recapitulate endogenous DP expressing NHEKs (Figure 3E-F). Conversely, B55α knockdown had no significant impact on the cell shapes in S2849G-DP-GFP expressing cells that are constitutively hypo-phosphorylated (Figure 3H, Supplemental Figure 4E-G). Together, these data suggest that PP2A-B55α regulates the cells mechano-response through phospho-regulation of DP’s C-terminus.

### PP2A-B55α is necessary to maintain epidermal adhesive capacity upon mechanical stretch

To understand the functional relevance of stretch-dependent B55α-mediated phospho-regulation of DP, an adhesion assay was performed. NHEK’s with and without B55α knockdown were subjected to 24 hours of cyclic stretch before keratinocyte monolayer adhesions were assessed via a dispase assay. Intact monolayers were lifted off culture dishes with dispase treatment (pre-stress) and subjected to mechanical rotations to induce monolayer fragmentation (post-stress) (Figure 3I, Supplemental Figure 4H). Scramble control monolayers remained intact regardless of whether they were subjected to stretching prior to the dispase assay (Figure 3J). Additionally, under these conditions, loss of B55α had no significant impact on the extent of monolayer fragmentation in unstretched controls. However, upon 24 hours of stretch, B55α knockdown significantly increased monolayer fragmentation by greater than 2-fold. These data suggest that PP2A-B55α regulation plays a key role in maintaining keratinocyte adhesive capacity when subjected to mechanical strain.

### Hypo-phosphorylated desmoplakin is enriched in the superficial layers of human epidermis

Our data support the existence of a cellular response to external mechanical stretch requiring the phospho-regulation of the DP-IF connection to maintain keratinocytes’ adhesive capacity. However, it remains unclear how keratinocytes use this regulatory pathway during normal homeostasis. Previous studies identified an epidermal tension gradient across the apical to basal aspect of the epidermis, with the highest tension in the superficial granular layers (Broussard et al., 2021; Fiore et al., 2020). To determine whether the DP-IF connection is phospho-regulated across the epidermal mechanical gradient, cross sections of human skin samples were collected and stained with antibodies recognizing total DP, single phospho-S2849 (pDP), or a double-phospho-DP (ppDP) antibody that recognizes DP phosphorylated at both the S2849 priming site and an upstream phospho site within the phospho-motif, S2845 (Figure 4A-E). Staining with either the pDP or ppDP alongside total DP revealed a phosphorylation gradient across the layered epidermis with phosphorylated DP significantly decreased in the upper layers (Figure 4F-G, 4J-K). Strikingly, the SG1 layer showed a near complete loss of pDP and ppDP, as quantified using line scans to measure pDP/ppDP signal at the membranes of each SG layer and the percent of membranes that were pDP/ppDP-positive (Figure 4G-I, 4K-M). These data support the presence of a phospho-DP gradient inversely associated with the previously reported tension gradient of the epidermis. Together, these data suggest that the skin actively regulates the DP-IF linkage through phosphorylation in coordination with the epidermal mechanical gradient, raising the possibility that DP de-phosphorylation both responds to and contributes to the spatial organization of mechanical forces across the tissue.

**Figure 4:**
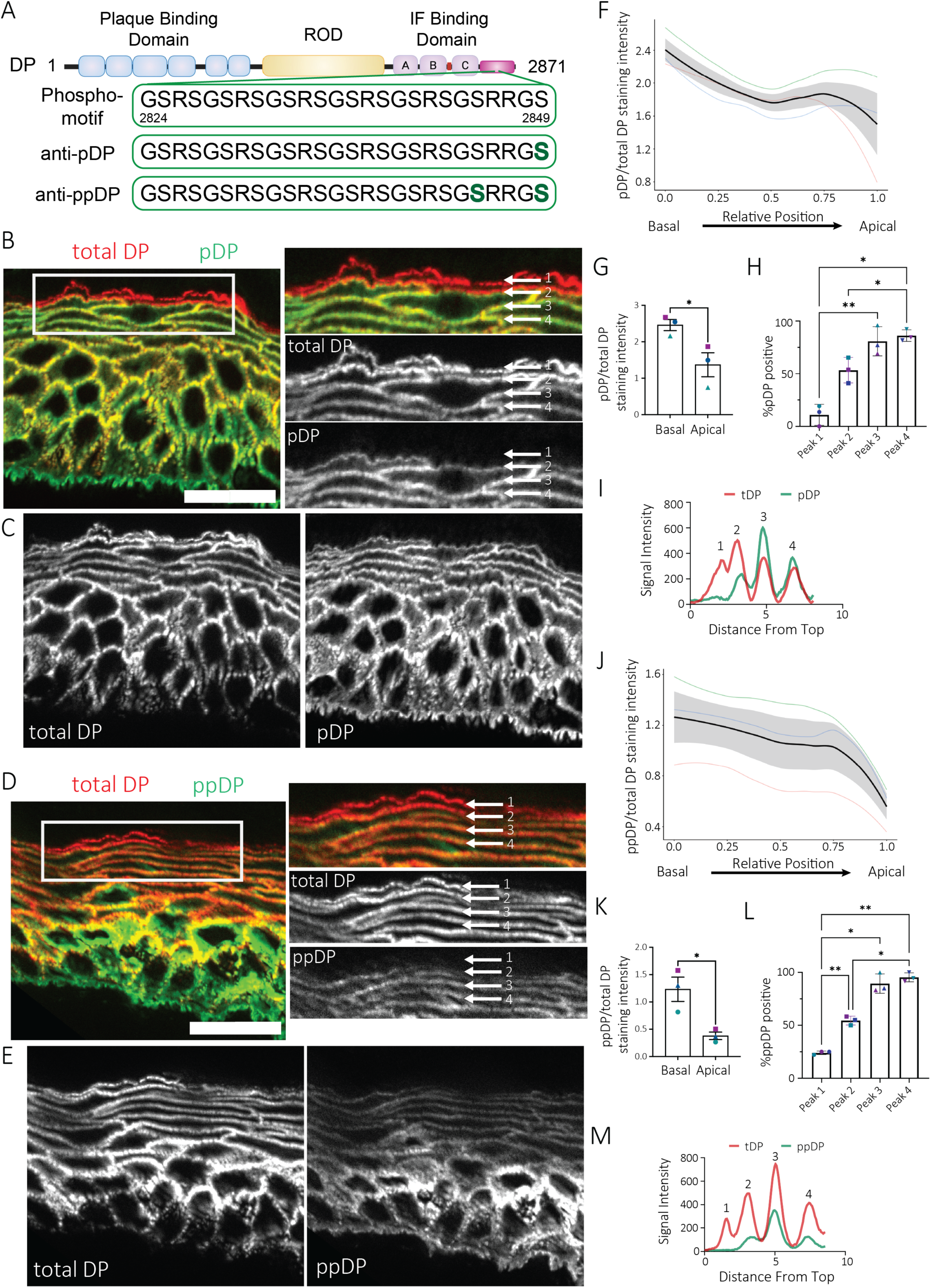
Dephosphorylated DP correlates with the high-tension layers of the human stratified epidermis. A) A schematic of DP visualizing the phosphor-regulatory motif, including the phospho-sites recognized by the pDP and ppDP antibodies. B-C) Cross sections of human skin samples stained with total DP (red) and phospho S2849 DP (pDP, green). D-E) Cross sections of human skin samples stained with total DP (red) and double-phospho-S2845/S2849 DP (ppDP, green). F) Trendline data from pDP/total DP staining intensity at cell membranes along the basal to apical aspect of the skin. Data was taken from linescans and the relative position across the sections was normalized to a 0-1 scale. Each colored line represents the trendline from an individual human sample. Black line represents the average Loess trendline across the n=3 with the gray ribbon representing the SD. G) Graph of the average p/tDP staining intensity at the basal (0 position) and apical (1 position) membranes in (F). H) Graph showing the percent of pDP positive membranes as determined by the percent of total DP peaks. I) A representative line scan measuring staining intensity in the granular layer for both total and pDP used in (H). Each peak represents a cell membrane of the SG1-SG3 layer cells. J) Trendline data from ppDP/total DP staining intensity at cell membranes along the basal to apical aspect of the skin. Data was taken from linescans and the relative position across the sections was normalized to a 0-1 scale. Each colored line represents the trendline from an individual human sample. Black line represents the average trendline across the n=3 with the gray ribbon representing the SD. K) Graph of the average pp/tDP staining intensity at the basal (0 position) and apical (1 position) membranes in (J). L) Graph showing the percent of ppDP positive membranes as determined by the percent of total DP peaks. M) A representative line scan measuring staining intensity in the granular layer for both total and ppDP used in (L). Each peak represents a cell membrane of the SG1-SG3 layer cells. Images were acquired with an apotome confocal. Statistical analyses are from a two-way Anova with multiple comparisons from an n=3. Scale bars are 20 μm. P-values are represented by (*) = 0.05, (**) = 0.01, (***) = 0.001, (****) = 0.0001.

### Carvajal-associated DP mutants are unable to respond to mechanical stretch

A number of epidermal disorders have been linked to dysregulation of the DP-IF connection. Carvajal syndrome is a cardio-cutaneous disorder associated with non-synonymous mutations within the *DSP* gene leading to palmar plantar keratoderma, woolly hair, and dilated cardiomyopathy (Boule et al., 2012). While Carvajal-associated mutations can be seen throughout *DSP*, one common feature is the loss of the C-terminal region of DP, including part of the IF binding domain and the previously characterized phospho-regulatory motif (Boule et al., 2012; Chalabreysse et al., 2011). Therefore, we were interested in identifying whether *DSP*-mediated skin disorders such as Carvajal have a dysregulated response to mechanical strain. To address this question, a rescue experiment was performed in the cr*DSP*-1 cells using WT-DP-GFP or two truncating forms of DP, one of which mimics a previously reported patient-associated Carvajal frameshift mutation harboring 17 extra amino acids, and the other truncating DP at the site of the missense mutation (Figure 5A)(Chalabreysse et al., 2011). The mutation, lying between plakin repeat B and C domains, results in a loss of the C-terminal portion of the IF binding domain and the entire phospho-regulatory domain (Figure 5A). Rescue with either Carvajal-DP-GFP construct resulted in impaired restoration of the stretch-induced cell elongation seen with WT-DP-GFP (Figure 5B-D, Supplemental Figure 5A-B).

**Figure 5:**
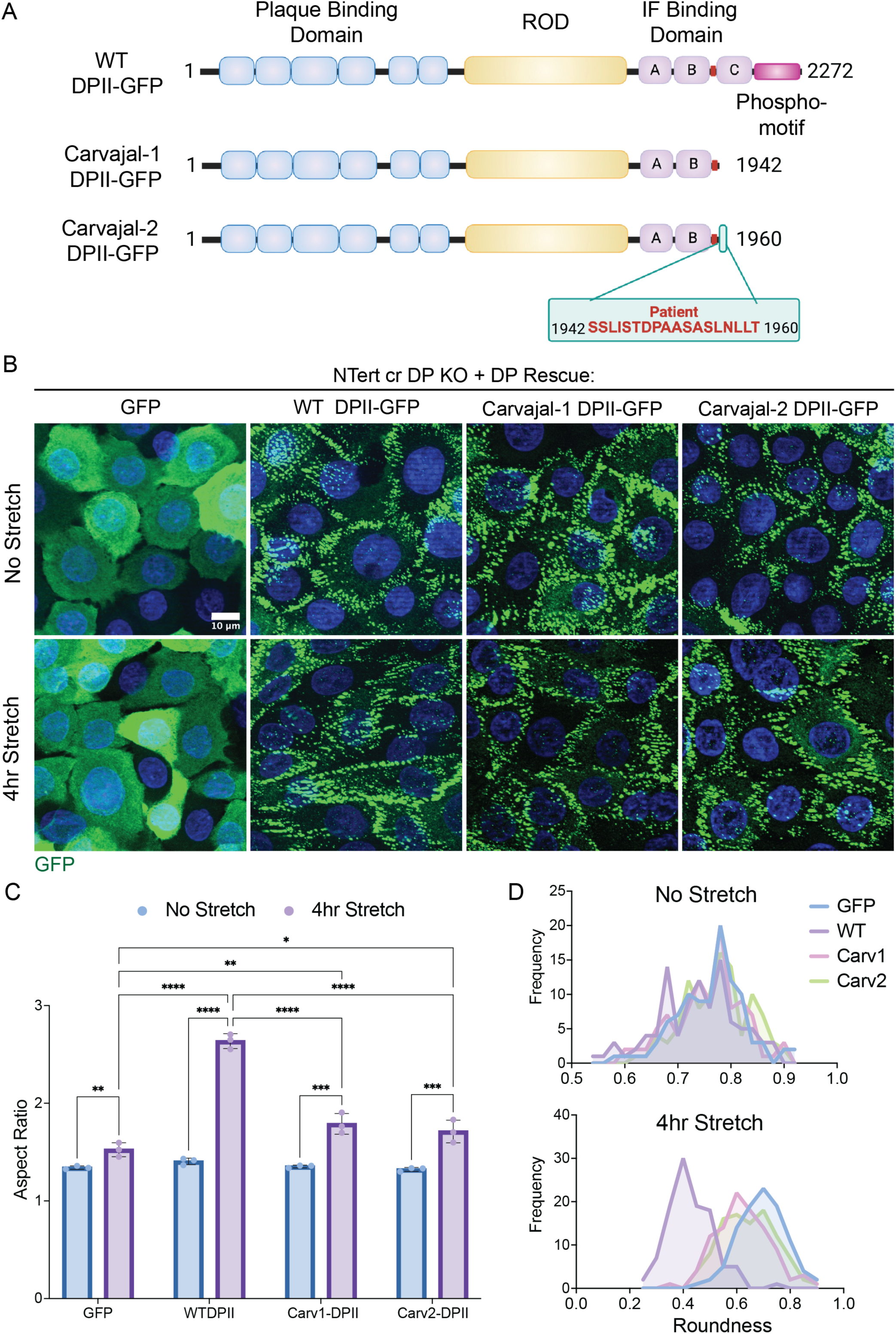
Carvajal patient-derived DP mutants are unable to adapt their cell shapes in response to mechanical stretch. A) A schematic of DPII visualizing the Carvajal-mutant constructs used, including a truncation mutant missing the deleted portion of DPII from the C-terminus (Carvajal-1) and a construct comprising the exact patient sequence including the addition of 18 amino acids onto DPII and an early stop codon (Carvajal-2). B) DP knockout cells cr*DSP*-1 were transduced with either WT-DPII GFP, Carvajal-1 DPII GFP, or Carvajal-2 DPII GFP and stretched for 4 hours. Representative images from no stretch and 4hr stretched conditions stained for GFP are shown. C) Graph of mean aspect ratio values from 3 biological replicates from (B). D) Frequency plots representing roundness measurements of individual cells from each condition in (B). Plots are separated by treatment. Statistical analyses are from a two-way Anova with multiple comparisons from an n=3. Scale bar is 10 μm. Images were acquired with an apotome confocal. P-values are represented by (*) = 0.05, (**) = 0.01, (***) = 0.001, (****) = 0.0001.

To understand how this impaired mechanical response impacts the epidermal structure in Carvajal syndrome, we examined the epidermis of a Carvajal mouse model containing a frameshift mutation at the end of DP’s plakin repeat B domain resulting in a premature truncation very similar to that of the human mutation analyzed above (Figure 6A)(Herbert Pratt et al., 2015). Earlier reports characterized these mice as having ruffled, woolly hair, epidermal abnormalities, and cardiac dysfunction in homozygous mice phenocopying Carvajal syndrome. Skin was excised from the dorsal side of wild-type and homozygous E18 mice and prepared as whole mounts to visualize the 3D structures of the epidermis (Figure 6B, Supplemental Movie 1-2). Initial observations revealed elongated cellular phenotypes in the superficial cell layers of wild-type mice compared to the basal layer (Figure 6C-D, Supplemental Movie 1-2). These polarized cell shapes were seemingly dampened in the homozygous Carvajal mice. To quantify the epidermal cell shapes in specific layers, a previously reported “cell shape index” was used to measure cells in cross sections of E18 mice stained with E-Cadherin and ZO-1 as markers of the cell membrane (Figure 6E). It was previously shown that the cell shape index is higher in granular layer cells compared to basal cells, consistent with a more elongated cell shape observed in the wild-type whole mount staining (Soffer et al., 2026). In contrast, cells in the granular layers of Carvajal mouse skin exhibited lower shape index values than wild-type mice (Figure 6F). The shape index values are consistent with other cell shape metrics such as AR values, which were also significantly lower in granular layer Carvajal cells (Figure 6G). Together, these data are consistent with the idea that Carvajal-associated mutations in DP have an impaired mechano-response to tension that may impact their ability to properly adapt as the cell experiences the epidermal mechanical gradient.

**Figure 6:**
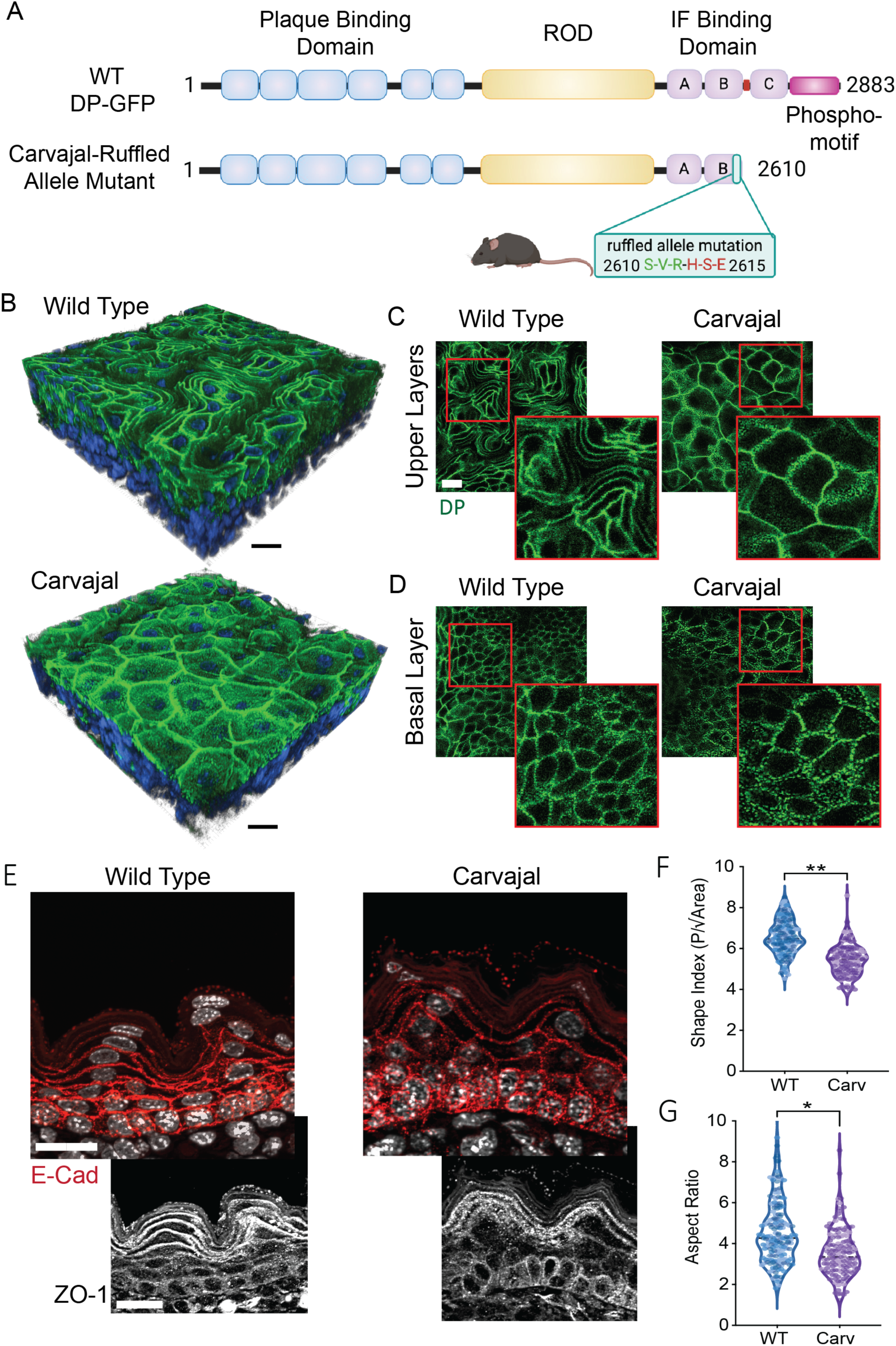
Carvajal mouse model exhibit aberrant cell shapes in vivo. *A) A schematic of* wildtype DP (top) compared to the DP mutation found in the ruffled mouse (bottom) showing the 3 unique amino-acid addition and truncation location of the DP mutation. B) 3D reconstructed images of whole mounts of mouse dorsal skin stained with DP (green) and DAPI (blue) from wildtype (top) and Carvajal (bottom) mice. Images were acquired using a multiphoton microscope. C-D) Apical view of a single Z-plane of whole mounts in (B) stained for DP. Images show z-slices corresponding to the upper stratified layers (C) and the apical surface of the basal layers (D). E) Cross sections of mouse dorsal skin from wildtype (left) and Carvajal (right) mice stained with E-Cadherin (red), DAPI (white), and ZO-1 (lower, white). Images were acquired with an apotome confocal. F) Shape index measurements from 3 biological replicates of (E). Points represent sign cell measurements, and varying color shades denoted different biological repeats. G) Aspect ratio measurements from 3 biological replicates of (E). Points represent sign cell measurements, and varying color shades denoted different biological repeats. Scale bars are 20 μm. Statistical analyses are from a two-way Anova with multiple comparisons from an n=3. P-values are represented by (*) = 0.05, (**) = 0.01, (***) = 0.001, (****) = 0.0001.

## DISCUSSION

As the foremost barrier between our body and the outside world, maintaining mechanical resilience is a fundamental requirement of the epidermis. To maintain a functional barrier, keratinocytes within the skin must respond and adapt to a constantly changing mechanical environment (Biggs et al., 2020; Mao and Wickstrom, 2024; Peskoller et al., 2022). Here we identified desmosomes, and specifically the DP-IF linkage, as a critical node in the epidermal mechano-response that is actively regulated in response to mechanical strain and is important for the normal homeostatic architecture of the stratified epidermis.

Prior work has focused on the role of the actin network in driving the epidermal mechano-response. In our study, while dysregulation of the DP-IF connection dramatically affected cell reorientation in response to stretch, actin filaments remained aligned in a stretch-dependent manner. Therefore, our work suggests that actin orientation alone is insufficient to elicit a complete mechano-response and that the DP-IF network is required alongside actin and its associated junctions to facilitate an adaptive response to mechanical stretch. This is consistent with several reports from our group and others that describe a collaborative role for the cytoskeletal networks that drive many critical processes (Broussard et al., 2021; Nanavati et al., 2024; Quinlan et al., 2017; Rubsam et al., 2023).

Our previous work demonstrated that the DP-IF connection can be tuned through modulation of a serine phosphorylation cascade within a C-terminal unstructured motif in DP (Stappenbeck et al., 1994). DP phosphorylation leads to a more dynamic interaction with IF whereas de-phosphorylation leads to a stronger, more stable interaction, associated with increased adhesive capacity and cell stiffness (Bartle et al., 2020; Broussard et al., 2017; Hobbs and Green, 2012). We now show that mechanical strain activates this mechanism, resulting in PP2A-B55α-dependent DP dephosphorylation necessary for the adaptive cell reorientation required to reestablish homeostasis. Furthermore, we provide evidence that this phospho-regulation helps maintain adhesive capacity specifically when epithelial sheets are subjected to stretch. Phospho-regulatory domains have been identified in other regions of DP that have been shown to impact cell adhesion, however it is unclear whether these other phospho-sites contribute to similar mechano-responsive properties (Rathod et al., 2024). In accordance with our results, an unbiased phospho-proteomics dataset from stretched keratinocytes revealed that the S2849-DP phospho-peptide decreased upon initial exposure to stretch but recovered overtime (Nava et al., 2020). Together these data are consistent with previous reports describing DP undergoes structural changes to bear mechanical load under acute conditions but not homeostatic conditions (Dong et al., 2025; Price et al., 2018; Sadhanasatish et al., 2023; Seeley et al., 2026). Importantly, these previous studies showed that the orientation of strain determined the mechanical load experienced by DP, and the constitutively hypo-phosphorylated DP experienced mechanical load. These observations are consistent with our data showing that hypo-phosphorylated DP is capable of eliciting a proper mechano-response. Overall, these data highlight the importance of understanding how DP post-translational regulation facilitates the cell’s adaptive response to mechanical strain.

Towards understanding how DP phospho-regulation regulates mechanics in vivo, we assessed changes in DP’s phospho-signature across the apical-basal aspect of the human stratified epidermis using antibodies requiring that either one (pDP) or two (ppDP) phospho-sites be modified. Considered together, these antibodies measure the extent to which the phosphorylation motif cascade in DP has been initiated. Given that hypo-phosphorylated DP results in increased DP-IF interactions associated with cell stiffness, we propose that this signature can provide higher resolution information about the previously identified gross differences in tension between basal and suprabasal epidermal layers. Using these antibodies, we observed a phospho-gradient across the stratified layers with a peak loss of DP phosphorylation in the granular layers, consistent with the idea that the tension gradient exists across the apical-basal aspect of the multi-layered epidermis (Broussard et al., 2021; Fiore et al., 2020). These data support the idea that the DP-IF connection is tuned by modulating the number of phosphorylation sites on DP to help the epidermis adjust to changes within the epidermal mechanical environment. However, more work is needed to understand the extent by which DP phospho-regulation contributes to the formation and maintenance of the epidermal tension gradient.

This DP phospho-gradient occurs in concert with gradients of actomyosin machinery and actin-spectrin networks required for morphological changes that occur as cells transit into superficial layers (Rubsam et al., 2017; Rubsam et al., 2023; Soffer et al., 2026). How DP phosphorylation and consequent increased DP-IF association are coordinated with progressive changes in the actin organization to support differentiation and barrier formation is unclear. We’ve previously observed that the superficially concentrated desmosomal cadherin, desmoglein 1, nucleates an actin remodeling complex, raising the possibility that differentiation-specific desmosome complexes could link these systems in a layer specific manner (Nekrasova et al., 2018). It is also unclear what drives the DP phosphorylation changes throughout the layers. Previous studies identified an apical-basal gradients of ErbB family kinases, including EGFR, that coordinates junctional functions required for establishing the epidermal barrier (Green et al., 2022; Rubsam et al., 2017; Soffer et al., 2026). Moreover, these same EGFR-dependent signaling pathways that direct granular layer function are shown to be activated upon intercellular stretch and capable of inhibiting GSK3β, the kinase responsible for phosphorylating DP’s S2849 site (Hino et al., 2020; Hirashima et al., 2023; Ma et al., 2007). Furthermore, staining of GSK3β in human skin shows an apical-basal gradient of GSK3β activity inversely correlated with our phospho-DP staining, with inactive GSK3β concentrated in the upper layers (Lee and Ro, 2015; Ma et al., 2007). Building on this data, it is possible that these or other differentiation-dependent signaling pathways can affect DP phosphorylation in a layer-specific manner. Understanding how the distribution and activation of DP’s upstream regulators, including GSK3β and PP2A-B55α are distributed across the apical-basal aspect will be important for better understanding how DP facilitates epidermal mechanical responses (Albrecht et al., 2015).

In addition to the phospho-gradient across the multi-layered epidermis, staining with both ppDP and pDP antibodies revealed an abrupt loss of signal at the SG1/SG2 transition, with 75-80% of cells in the SG1 layer having a complete loss of phosphorylated-DP. This finding raises the question: why is the SG1 uniquely hypo-phosphorylated? Several unique characteristics of the SG1 layer could contribute to the observed pattern. As the layer above the tight junctions, desmosomes are the only cell-cell adhesion junction present in SG1 (Peskoller et al., 2022; Yoshida et al., 2013). Additionally, the SG1 layer has the highest prevalence of cortical actin, a driver of intercellular tension (Rubsam et al., 2017). Thus, it is possible that in the SG1 layer DP is forced to bear more mechanical strain and therefore requires a tighter DP-IF connection, which in turn requires a fully hypo-phosphorylated state. Further, as the last living layer of the skin, the SG1 layer is engaged in a cross-linking process required to transition into the cornified layer (Fuchs, 2007; Peskoller et al., 2022). The cross-linking process may interrupt the mechanical and/or enzymatic activity keeping DP in a phosphorylated state. More studies are necessary to determine why SG1 cells induce a near complete dephosphorylation of DP’s C-terminus, but our in vitro data raise the strong possibility that the epidermal mechanical environment in the SG cells is a contributing factor.

Our study also raises the possibility that DP-associated disease phenotypes may result at least in part from a failure of cells to adapt to their mechanical environments. We show that in Carvajal syndrome, dysregulation of the DP-IF connection leads to an impaired mechano-response in cells and identified a disorganized structure within Carvajal epidermis. Altogether, our data suggest an association between Carvajal-mutant cells’ ability to regulate its DP-IF connection in response their mechanical environment and their epidermal structure. But several questions remain. Firstly, it is unclear the extent to which the Carvajal-associated phenotype is driven by a disruption to the IF binding site, or a lack of a phospho-regulatory motif to allow for modulation of the DP-IF binding affinity. More clear is that dysregulation of the DP-IF linkage in Carvajal syndrome impairs the epidermis’ mechano-response. How this impaired mechano-response relates to the disease pathology remains elusive. Future studies are required to better understand the contributions of mechanical response pathways to Carvajal pathology. These include understanding whether the desmosome-mediated mechano-response contributes to the epidermal thickening, as well as its role in the localized presentation of the pathology in the highly mechanically stressed sites of the palmar plantar epidermis (Boule et al., 2012; Chalabreysse et al., 2011; Protonotarios and Tsatsopoulou, 2004). Further elucidating the role of mechano-response pathways in the context of Carvajal could provide a better understanding of the impact of mechanical adaptation in other Epidermal Differentiation Disorders (EDD’s) including other palmar plantar keratodermas or diseases affecting the DP-IF network (Hernandez-Martin et al., 2025; Paller et al., 2025).

## MATERIAL AND METHODS

### Cell lines and culture condition

TERT immortalized keratinocytes, NTert-2G (Dickson et al., 2000), cells were cultured in Keratinocyte SFM media with supplements (Gibco), 0.3 mM CaCl^2^, and 1% penicillin/streptomycin. Normal Human Epidermal Keratinocytes (NHEKs) were isolated from neonatal foreskin provided by the Northwestern University Skin Biology and Disease Resource-based Center (SBDRC) as previously described (Arnette et al., 2016). NHEKs were maintained in M154 (Thermo Fisher Scientific) growth media supplemented with 0.07 mM CaCl^2^, human keratinocyte growth supplement (HKGS) (Thermo Fisher Scientific), gentamicin and amphotericin B. Cell lines were maintained at 37°C in a humidified atmosphere of 5% Co^2^. Media supplemented with high Ca^2+^ (1.2mM for NHEKs, 1.3 mM for NTerts) for experimental treatments as described.

For siRNA transfection, an Amaxa Nucleofector System was used according to the manufacturer’s instructions. Cells were suspended in Ingenio Electroporation Solution (Mirus Bio) with Dharmacon ON-TARGET siRNA (Horizon Discovery) targeting the B55α gene, PPP2R2A (J-004824-06) at a final concentration of 50 mM and electroporated using program X-001. Treated cells were plated at confluency on culture dishes.

### Crispr cell line generation and sequencing

The Alt-R CRISPR-Cas9 system (Integrated DNA Technologies) and predesigned guideRNA targeting the *DSP* gene (Hs.Cas9.DSP.1.AC) or a negative control guideRNA (CGTTAATCGCGTATAATACG) was used to generate cell lines. Cas9 and gRNA were transfected into NTert cells using the Amaxa Nucleofector System. RNAs were resuspended in IDTE buffer at a 200uM concentration and the gRNA and tracrRNA were mixed to a final concentration of 44 uM before incubation at 95°C for 5 minutes. A 1:1 mixture of gRNA:Cas9 was incubated for 20 minutes at room temperature prior to being added to cell suspension in Ingenio Electroporation Solution (Mirus Bio). The final mixture was electroporated using program X-001 and cells were grown and split into single cell colonies. Colonies were isolated and propagated into clonal cell lines.

### Viral expression protocol

Retrovirus was generated using the Phoenix amphotropic cell line (Gary Nolan, Stanford University Medical School) maintained in DMEM (Thermo Fisher Scientific) with 10% FBS and antibiotics. Lipofectamine reagent was used to transfect Phoenix cells with retroviral constructs, which were selected with 1 mg/ml puromycin in culture medium. Retroviral supernatants were harvested from Phoenix cells cultured at 32°C and concentrated on a Centricon Plus-20 column (MilliporeSigma). Subconfluent NHEK cultures were incubated at 32°C in M154 (Thermo Fisher Scientific) along with 4 μg/ml polybrene (Sigma Chemical Co.) and concentrated virus for 2–4 hours. Viral media was washed and cells were cultured under normal maintenance conditions.

### Immunofluorescence and microscopy

Cells were fixed with 4% paraformaldehyde (PFA) solution for 20 minutes at room temperature followed by permeabilization with either anhydrous ice-cold methanol for 2 minutes at −20°C or 0.25% Triton X PBS on ice for 15 minutes to visualize actin filaments. Cells were blocked with 5% goat serum in PBS for 1 hour at 37°C. Phalloidin staining was done for 1 hour at 37°C. Glass coverslips were mounted on samples with ProLong Gold anti fade reagent (Thermo Fisher Scientific) and allowed to cure for 48 hours at room temperature.

Human skin samples were stained from frozen OCT embedded samples. Slides were fixed with anhydrous ice-cold methanol for 3 minutes at −20°C. Samples were blocked with 5% goat serum for 1 hour at 37°C and incubated overnight at 4°C with primary antibodies in a humidified chamber followed by PBS washes and secondary antibody incubation for 1 hour at 37°C. Glass coverslips were mounted on samples with ProLong Gold anti fade reagent (Thermo Fisher Scientific) and allowed to cure for 48 hours.

Mouse sections were stained from paraffin embedded samples. Embryo samples were fixed whole with 10% neutral-buffered formalin, embedded in paraffin, and cut into 4-to 5-μm sections. Paraffin-embedded sections were baked at 60°C overnight and deparaffinized using xylene. Samples were run through a series of ethanol and PBS dips, and slides were permeabilized in 0.5% Triton X. Antigen retrieval was performed by heating samples to 95°C in 1.8mM citric acid and 8.2mM sodium citrate buffer for 20min, as described in (Mescher et al., 2017). Sections were blocked in 1%BSA with 2% goat serum for 1 h at 37°C and incubated with primary antibodies overnight at 4°C in a humid chamber and secondary antibodies at 37°C for 1 h followed by mounting using ProLong Gold (Thermo Fisher Scientific).

Images were acquired using ZEN 2.3 software with an epifluorescence microscope system (Axio Imager Z2, Carl Zeiss) fitted with an X-Cite 120 LED Boost System, an Apotome.2 slide module, Axiocam 503 Mono digital camera, and a Plan-Apochromat 40x/1.4 or 60x/1.4 objectives (Carl Zeiss). Actin and keratin images were acquired on a Nikon Perfect Focus Ti2 inverted spinning disk microscope equipped with a Yokogawa CSU-W1 SoRa system, a Hamatsu Orca-Fusion Camera, and a 60x Plan-Apochromat Lambda D lens 1.42 NA (Nikon). Images are processed using FIJI ImageJ software (Schindelin et al., 2012).

### Whole mount imaging

Protocol was adapted from (Cetera et al., 2024). E18.5 embryos were harvested and placed into PBS with magnesium and calcium. Dorsal skin was dissected off each embryo using micro dissecting scissors, non-serrated forceps, and pipettes tips affixed with a single toothbrush bristle, which was used to sweep under the skin and break apart connective tissue. Pelts were fixed for 1.5hrs at 22°C while rocking in 4% PFA. Pelt was cut into multiple pieces before incubation in 5% NGS/0.5% Triton-X permeabilizing solution overnight at 4°C. After blocking, samples were washed with PBS then incubated overnight in primary antibodies at 4°C and the same was done for secondary antibody incubation. After secondary, samples were washed for 3x 30min PBS washes before mounting epidermis down in a 24-well glass bottom dish using Gelvatol. A coverslip was placed on top of the tissue and gently pressed down to keep the tissue flat for imaging. Whole mounts were imaged using a Leica DiveB Sp8 Multiphoton microscope with a 25x water immersion objective.

### Western blot analysis

Whole cell lysates were collected using Urea Sample Buffer. Proteins were separated by SDS-page electrophoresis on 4-15% Mini-PROTEAN TGX Gradient gels (BioRad) and transferred in a submerged wet transfer overnight at 4 mV onto a 0.2 nitrocellulose membrane. Primary antibodies were incubated overnight at 4°C and secondary antibodies were incubated for 1 hour at 22°C. Proteins were visualized using a LI-COR OdysseyXF.

### Antibodies and reagents

The following primary antibodies were used: NW161 rabbit anti-DP N-terminal (Bornslaeger et al., 1996); 11-5F mouse anti-DP c-terminal (Sigma, Gift from D. Garrod)(Parrish et al., 1987); anti-phospho-S2849 DP (Albrecht et al., 2015; Bouameur et al., 2013); anti-dual phospho-S2845/S2849 DP (Albrecht et al., 2015; Perl et al., 2023); 1407 chicken anti-Pkg (Aves Laboratories); 2G9 Mouse anti-B55α (Cell Signaling Technology); Mouse anti-E-cadherin (BD Transduction, BD610181); Rabbit anti-ZO-1 (Invitrogen 40-2200); mouse anti-GAPDH (Santa Cruz Biotechnology); hVinc1 mouse anti-vinculin (Millipore Sigma); chicken anti-GFP (Thermo Scientific); JL-8 mouse anti-GFP (Clontech); rabbit anti-GFP (Clontech). Alexa Fluor 568 Phalloidin (Thermo Fisher Scientific).

The following secondary antibodies were used: goat anti-mouse IgG HRP (Cell Signaling Technologies); goat anti-rabbit IgG HRP (Cell Signaling Technologies); Goat anti-Mouse conjugated with Alexa Fluor-488/568 (Thermo Fisher Scientific); Goat anti-Chicken conjugated with Alexa Fluor-647 (Thermo Fisher Scientific); Goat anti-Rabbit conjugated with Alexa Fluor-488/568 (Thermo Fisher Scientific).

The WT-DPII-GFP construct and adenovirus generation using Gateway recombination were previously described whereby DPII-GFP was cloned into the pAd CMV/V5-DEST vector or the LZRS retroviral backbone (Mitchell Denning; Loyola University Medical School)(Kam et al., 2018). The DPNTP-GFP (Broussard et al., 2021), S2849G-DPII-GFP, the Carv1-DPII-GFP and Carv2-DPII-GFP (Zarkoob et al., 2026) constructs were created by generating a point mutation in the WT-DPII-GFP constructs (Epoch Life Science).

### Cell stretching

Cells were plated at confluency on Collagen I coated BioFlex 6-Well culture plates (Flexcell International Corporation) in normal growth media and put into high Ca^2+^ media for 24 (NHEKs) or 48 (NTerts) hours prior to stretching. BioFlex plates were put on the FlexCell FX-6000 Tension System (Flexcell International Corporation) with equibiaxial posts and grease was applied to suction posts prior to loading. A cyclic stretching regimen of 18% strain in 7 second cycles was performed for the duration of the treatment as noted.

### Dispase Assay

Cells were stretched as described above for 24 hours prior to being washed in PBS and treated with 2.4 U/mL of dispase (Sigma Millipore) diluted in PBS containing Ca^2+^ for 30 min. Lifted monolayers were placed in 15mL conical tubes containing 3 mL PBS and inverted 10-15 times as described in (Hudson et al., 2004). Resulting fragments were returned to a fresh 6-well dish and imaged using a dissecting microscope (MZ6; Leica).

### Mouse handling

All housing care and use of animals was handled according to Northwestern University Institutional Animal Care and Use Committee (IACUC) Protocols (ID IS00017230). Carvajal mice originally reported by (Herbert Pratt et al., 2015) were obtained from Jax (Strain ID: 005362) via cry recovery on a C57BL/6J background and maintained as heterozygotes. Mice were housed in a barrier facility in a temperature-controlled room with a 12-hour light cycle and given ad libitum access to food and water. For timed matings, mice were mated overnight and separated the following morning to generate timed pregnancies.

### Image quantification

All image quantification was conducted using FIJI ImageJ (Schindelin et al., 2012). For 2D cultures, basal cell layers were used for quantification to control for differences in stratification between conditions. AR, Feret values, roundness, and cell orientation data were acquired by cell border tracing of using membrane markers as noted in figure legends. Line scans across cell borders were used to calculate membrane staining intensity values in Figure 3. Values are presented as a ratio of phosphoDP/total DP intensity. Anisotropy measurements were done using the FribrilTool ImageJ plugin tool (Boudaoud et al., 2014).

Quantification of tissue samples were performed on cross sections. In Figure 4, line scans were drawn from above epidermal tissue capturing signal intensity across the granular layer. Total DP stain was used to identify peaks representing cell borders, and peaks were overlayed with phospho-DP intensity values to identify corresponding pDP signal. A loess model was used to calculated overall trendlines from data. %pDP/ppDP positive membranes were calculated using the number of total DP peaks that corresponded with a phospho-DP peak. In Figure 6, cells shape analysis was conducted on ZO-1 positive cell layers based on E-Cad membrane staining. DAPI was used to identify whole cells in the cross section. A cell shape index (perimeter (P)/√Area) was applied using measurements acquired by ImageJ (Soffer et al., 2026).

### Statistical analysis and data plotting

Plots were generated using GraphPad Prism 10, R Studio, and Adobe Illustrator. All statistics were performed on raw data, although some plots display fold change for visual clarity. Significances were determined via a paired one-way analysis of variance (ANOVA) with multiple comparisons, except where indicated. Statistics conducted on frequency plots adjusted to a 360° scale.

## DATA AVAILABILITY

No large datasets were generated or analyzed during the current study. All data generated or analyzed during this study are included in this published article and its supplementary information files. The datasets used and/or analyzed during the current study are available from the corresponding author upon reasonable request.

## ACKNOWLEDGEMENTS

We thank members of the Green lab for their valuable feedback and discussion. We also thank Matthias Rübsam for his guidance with utilizing the cell shape index analysis. Research reported in this publication was supported by Northwestern University Skin Biology & Diseases Resource-Based Center of the National Institutes of Health under award number P30AR075049. Imaging work and the FlexCell stretching was performed at the Northwestern University Center for Advanced Microscopy (RRID: SCR_020996) generously supported by CCSG P30 CA060553 awarded to the Robert H Lurie Comprehensive Cancer Center. Confocal microscopy was performed on a Nikon SoRa system, purchased through the support of NIH 1S10OD032270-01. Northwestern University’s Pathology Core Facility (PCF) and Mouse Histology and Phenotyping Laboratory (MHPL) performed the sectioning of human and mouse skin samples. This work was supported by NIH R01 AR43380, NIAMS R01 AR041836, and NCI R01 CA228196 and the JL Mayberry Endowment to KJG. A.L.P was supported by T32AR060710 and F32AR081677.

## DECLARATION OF GENERATIVE ARTIFICIAL INTELLIGENCE (AI) OR LARGE LANGUAGE MODELS (LLMs)

The author (s) did not use AI/LLM in any part of the research process and/or manuscript preparation.

## SUPPLEMENTAL FIGURE LEGENDS

***<u>Supplemental Figure 1:</u>*** A) Sanger sequencing of DP Crispr knockout cells generated NTERT-2G cells. 2 crControl clones were generated using a scramble guideRNA alongside 2 cr*DSP* clones generated with a guideRNA targeting exon 1 of the *DSP* gene. A 481 PCR product including the *DSP* guide RNA site was sequenced. B) Number of mismatches were calculated for each clone. C) Agarose gel visualizing the PCR product sequenced in (A). D) Immunoblot confirmation of DP expression in the 4 Crispr treated clones. DP C-terminus targeting antibody (115F) was used.

***<u>Supplemental Figure 2:</u>*** A) DP knockout cells were stretched for 4 and 24 hours. Immunofluorescence images of crControl-2 (top) and cr*DSP*-2 (bottom) cells post treatment detecting Pkg and DP. Images were acquired with an apotome confocal. Representative cells are outlined in red and reconstructed below image. Aspect ratio (AR) values of the representative cell are depicted below. Scale bars are 20 μm. B) Frequency plots representing cell orientation (Angle) of individual cells from each condition in (A). Plots are separated by treatment. C) Mean feret values from 3 biological replicates of (A). D) Rose plots of cell orientation spanning 0°-180° of NHEKs transduced with GFP or DPNTP that were stretched for 24 hours. Rho values for each rose plot are denoted below. Statistical analyses are from a two-way Anova with multiple comparisons from an n=3. P-values are represented by (*) = 0.05, (**) = 0.01, (***) = 0.001, (****) = 0.0001.

***<u>Supplemental Figure 3:</u>*** A) DP knockout cells cr*DSP*-1 were transduced with either GFP, WT-DPII GFP, or S2849G-DPII GFP. Representative images of cells after 0 or 4 hours of stretch and stained for GFP are shown. B) Immunoblot visualizing GFP expression in cell lysate of cells used in (A). C) Representative images of cells in (A) including crControl-1 cells after 0 or 4 hours of stretch and stained for Pkg are shown. D) Graph of mean aspect ratio values from 3 biological replicates from (B). E) Graph of mean feret values from 3 biological replicates from (B). F) Graph of individual cell aspect ratio values from (B, D). Varying color shades denoted different biological repeats. G) Frequency plots representing roundness measurements of individual cells from each condition in (B). Plots are separated by treatment. Images were acquired with an apotome confocal. Statistical analyses are from a two-way Anova with multiple comparisons from an n=3. Scale bars are 20 μm. P-values are represented by (*) = 0.05, (**) = 0.01, (***) = 0.001, (****) = 0.0001.

***<u>Supplemental Figure 4:</u>*** A) Graph of mean feret values from 3 biological replicates from NHEK cells with B55α knockdown via targeted siRNA were stretched for 4 hours. B) Graph of individual cell aspect ratio values from NHEK cells with B55α knockdown via targeted siRNA were stretched for 4 hours. C) Frequency plots representing roundness measurements of individual NHEK cells with B55α knockdown via targeted siRNA were stretched for 4 hours. Plots are separated by treatment. D) Immunoblot visualizing of B55α expression in NHEK cells with B55α knockdown via targeted siRNA. E) DP knockout cells cr*DSP*-1 were transduced with either WT-DPII GFP or S2849G-DPII GFP and B55α was knockdown in both conditions via targeted siRNA. Representative images of unstretched cells and stained for GFP are shown. F) Graph of mean feret values from 3 biological replicates from (E and Figure 3G). G) Graph of individual cell aspect ratio values from (E, Figure 3G-H). H) NHEK cells with B55α knockdown via targeted siRNA were stretched for 24 hours and treated with dispase to lift cell sheets and subjected to mechanical stress. Images of cell sheets prior to dispase assay mechanical stress. Scale bars are 20 μm. Images were acquired with an apotome confocal. Statistical analyses are from a two-way Anova with multiple comparisons from an n=3. P-values are represented by (*) = 0.05, (**) = 0.01, (***) = 0.001, (****) = 0.0001.

***<u>Supplemental Figure 5:</u>*** A) Graph of individual cell aspect ratio values from DP knockout cells cr*DSP*-1 transduced with either WT-DPII GFP, Carvajal-1 DPII GFP, or Carvajal-2 DPII GFP and stretched for 4 hours. B) Graph of mean feret values from 3 biological replicates from DP knockout cells cr*DSP*-1 transduced with either WT-DPII GFP, Carvajal-1 DPII GFP, or Carvajal-2 DPII GFP and stretched for 4 hours. Statistical analyses are from a two-way Anova with multiple comparisons from an n=3. P-values are represented by (*) = 0.05, (**) = 0.01, (***) = 0.001, (****) = 0.0001.

***<u>Supplemental Movie 1:</u>*** Movies of 3D reconstructed whole mounts from wildtype mouse dorsal skin. Movie on the left is stained with DP (green) and DAPI (blue) and movie on the right is stained with DP (green), actin (red), and DAPI (blue). Images were acquired using a multiphoton microscope.

***<u>Supplemental Movie 2:</u>*** Movies of 3D reconstructed whole mounts from homozygous Carvajal mouse dorsal skin. Movie on the left is stained with DP (green) and DAPI (blue) and movie on the right is stained with DP (green), actin (red), and DAPI (blue). Images were acquired using a multiphoton microscope.

## References

Albrecht LV, Zhang L, Shabanowitz J, Purevjav E, Towbin JA, Hunt DF, et al. GSK3-and PRMT-1-dependent modifications of desmoplakin control desmoplakin-cytoskeleton dynamics. J Cell Biol 2015;208(5):597–612. 10.1083/jcb.201406020.

Arnette C, Koetsier JL, Hoover P, Getsios S, Green KJ. In Vitro Model of the Epidermis: Connecting Protein Function to 3D Structure. Methods Enzymol 2016;569:287–308. 10.1016/bs.mie.2015.07.015.

Bartle EI, Rao TC, Beggs RR, Dean WF, Urner TM, Kowalczyk AP, et al. Protein exchange is reduced in calcium-independent epithelial junctions. J Cell Biol 2020;219(6). 10.1083/jcb.201906153.

Biggs LC, Kim CS, Miroshnikova YA, Wickstrom SA. Mechanical Forces in the Skin: Roles in Tissue Architecture, Stability, and Function. J Invest Dermatol 2020;140(2):284–90. 10.1016/j.jid.2019.06.137.

Bornslaeger EA, Corcoran CM, Stappenbeck TS, Green KJ. Breaking the connection: displacement of the desmosomal plaque protein desmoplakin from cell-cell interfaces disrupts anchorage of intermediate filament bundles and alters intercellular junction assembly. J Cell Biol 1996;134(4):985–1001. 10.1083/jcb.134.4.985.

Bouameur JE, Schneider Y, Begre N, Hobbs RP, Lingasamy P, Fontao L, et al. Phosphorylation of serine 4,642 in the C-terminus of plectin by MNK2 and PKA modulates its interaction with intermediate filaments. J Cell Sci 2013;126(Pt 18):4195–207. 10.1242/jcs.127779.

Boudaoud A, Burian A, Borowska-Wykret D, Uyttewaal M, Wrzalik R, Kwiatkowska D, et al. FibrilTool, an ImageJ plug-in to quantify fibrillar structures in raw microscopy images. Nat Protoc 2014;9(2):457–63. 10.1038/nprot.2014.024.

Boule S, Fressart V, Laux D, Mallet A, Simon F, de Groote P, et al. Expanding the phenotype associated with a desmoplakin dominant mutation: Carvajal/Naxos syndrome associated with leukonychia and oligodontia. Int J Cardiol 2012;161(1):50–2. 10.1016/j.ijcard.2012.06.068.

Broussard JA, Jaiganesh A, Zarkoob H, Conway DE, Dunn AR, Espinosa HD, et al. Scaling up single-cell mechanics to multicellular tissues - the role of the intermediate filament-desmosome network. J Cell Sci 2020;133(6). 10.1242/jcs.228031.

Broussard JA, Koetsier JL, Hegazy M, Green KJ. Desmosomes polarize and integrate chemical and mechanical signaling to govern epidermal tissue form and function. Curr Biol 2021;31(15):3275–91 e5. 10.1016/j.cub.2021.05.021.

Broussard JA, Yang R, Huang C, Nathamgari SSP, Beese AM, Godsel LM, et al. The desmoplakin-intermediate filament linkage regulates cell mechanics. Mol Biol Cell 2017;28(23):3156–64. 10.1091/mbc.E16-07-0520.

Cetera M, Sharan R, Hayward-Lara G, Devenport D. Evaluating Planar Cell Polarity in the Developing Mouse Epidermis. Methods Mol Biol 2024;2805:187–201. 10.1007/978-1-0716-3854-5_13.

Chalabreysse L, Senni F, Bruyere P, Aime B, Ollagnier C, Bozio A, et al. A new hypo/oligodontia syndrome: Carvajal/Naxos syndrome secondary to desmoplakin-dominant mutations. J Dent Res 2011;90(1):58–64. 10.1177/0022034510383984.

De R, Zemel A, Safran SA. Do cells sense stress or strain? Measurement of cellular orientation can provide a clue. Biophys J 2008;94(5):L29–31. 10.1529/biophysj.107.126060.

Dehner C, Rotzer V, Waschke J, Spindler V. A desmoplakin point mutation with enhanced keratin association ameliorates pemphigus vulgaris autoantibody-mediated loss of cell cohesion. Am J Pathol 2014;184(9):2528–36. 10.1016/j.ajpath.2014.05.016.

Dickson MA, Hahn WC, Ino Y, Ronfard V, Wu JY, Weinberg RA, et al. Human keratinocytes that express hTERT and also bypass a p16(INK4a)-enforced mechanism that limits life span become immortal yet retain normal growth and differentiation characteristics. Mol Cell Biol 2000;20(4):1436–47. 10.1128/MCB.20.4.1436-1447.2000.

Dong Y, Elgerbi A, Xie B, Han Y, Kwiatkowski AV, Choy JS, et al. Actomyosin forces trigger a conformational change in desmoplakin within desmosomes. Nat Commun 2025;16(1):9052. 10.1038/s41467-025-64124-4.

Fiore VF, Krajnc M, Quiroz FG, Levorse J, Pasolli HA, Shvartsman SY, et al. Mechanics of a multilayer epithelium instruct tumour architecture and function. Nature 2020;585(7825):433–9. 10.1038/s41586-020-2695-9.

Fuchs E. Scratching the surface of skin development. Nature 2007;445(7130):834–42. 10.1038/nature05659.

Fudge D, Russell D, Beriault D, Moore W, Lane EB, Vogl AW. The intermediate filament network in cultured human keratinocytes is remarkably extensible and resilient. PLoS One 2008;3(6):e2327. 10.1371/journal.pone.0002327.

Godsel LM, Hsieh SN, Amargo EV, Bass AE, Pascoe-McGillicuddy LT, Huen AC, et al. Desmoplakin assembly dynamics in four dimensions: multiple phases differentially regulated by intermediate filaments and actin. J Cell Biol 2005;171(6):1045–59. 10.1083/jcb.200510038.

Green KJ, Niessen CM, Rubsam M, Perez White BE, Broussard JA. The Desmosome-Keratin Scaffold Integrates ErbB Family and Mechanical Signaling to Polarize Epidermal Structure and Function. Front Cell Dev Biol 2022;10:903696. 10.3389/fcell.2022.903696.

Hatzfeld M, Keil R, Magin TM. Desmosomes and Intermediate Filaments: Their Consequences for Tissue Mechanics. Cold Spring Harb Perspect Biol 2017;9(6). 10.1101/cshperspect.a029157.

Herbert Pratt C, Potter CS, Fairfield H, Reinholdt LG, Bergstrom DE, Harris BS, et al. Dsp rul: a spontaneous mouse mutation in desmoplakin as a model of Carvajal-Huerta syndrome. Exp Mol Pathol 2015;98(2):164–72. 10.1016/j.yexmp.2015.01.015.

Hernandez-Martin A, Paller AS, Sprecher E, Akiyama M, Granier Tournier C, Aldwin-Easton M, et al. A proposal for a new pathogenesis-guided classification for inherited epidermal differentiation disorders. Br J Dermatol 2025;193(3):544–8. 10.1093/bjd/ljaf065.

Hino N, Rossetti L, Marin-Llaurado A, Aoki K, Trepat X, Matsuda M, et al. ERK-Mediated Mechanochemical Waves Direct Collective Cell Polarization. Dev Cell 2020;53(6):646–60 e8. 10.1016/j.devcel.2020.05.011.

Hirashima T, Hino N, Aoki K, Matsuda M. Stretching the limits of extracellular signal-related kinase (ERK) signaling - Cell mechanosensing to ERK activation. Curr Opin Cell Biol 2023;84:102217. 10.1016/j.ceb.2023.102217.

Hobbs RP, Green KJ. Desmoplakin regulates desmosome hyperadhesion. J Invest Dermatol 2012;132(2):482–5. 10.1038/jid.2011.318.

Hsu CK, Lin HH, Harn HI, Hughes MW, Tang MJ, Yang CC. Mechanical forces in skin disorders. J Dermatol Sci 2018;90(3):232–40. 10.1016/j.jdermsci.2018.03.004.

Hudson TY, Fontao L, Godsel LM, Choi HJ, Huen AC, Borradori L, et al. In vitro methods for investigating desmoplakin-intermediate filament interactions and their role in adhesive strength. Methods Cell Biol 2004;78:757–86. 10.1016/s0091-679x(04)78026-7.

Huen AC, Park JK, Godsel LM, Chen X, Bannon LJ, Amargo EV, et al. Intermediate filament-membrane attachments function synergistically with actin-dependent contacts to regulate intercellular adhesive strength. J Cell Biol 2002;159(6):1005–17. 10.1083/jcb.200206098.

Jin X, Rosenbohm J, Kim E, Esfahani AM, Seiffert-Sinha K, Wahl JK, 3rd, et al. Modulation of Mechanical Stress Mitigates Anti-Dsg3 Antibody-Induced Dissociation of Cell-Cell Adhesion. Adv Biol (Weinh) 2021;5(1):e2000159. 10.1002/adbi.202000159.

Kam CY, Dubash AD, Magistrati E, Polo S, Satchell KJF, Sheikh F, et al. Desmoplakin maintains gap junctions by inhibiting Ras/MAPK and lysosomal degradation of connexin-43. J Cell Biol 2018;217(9):3219–35. 10.1083/jcb.201710161.

Le HQ, Ghatak S, Yeung CY, Tellkamp F, Gunschmann C, Dieterich C, et al. Mechanical regulation of transcription controls Polycomb-mediated gene silencing during lineage commitment. Nat Cell Biol 2016;18(8):864–75. 10.1038/ncb3387.

Lee H, Ro JY. Differential expression of GSK3beta and pS9GSK3beta in normal human tissues: can pS9GSK3beta be an epithelial marker? Int J Clin Exp Pathol 2015;8(4):4064–73.

Lien JC, Wang YL. Cyclic stretching-induced epithelial cell reorientation is driven by microtubule-modulated transverse extension during the relaxation phase. Sci Rep 2021;11(1):14803. 10.1038/s41598-021-93987-y.

Ma C, Wang J, Gao Y, Gao TW, Chen G, Bower KA, et al. The role of glycogen synthase kinase 3beta in the transformation of epidermal cells. Cancer Res 2007;67(16):7756–64. 10.1158/0008-5472.CAN-06-4665.

Mao Y, Wickstrom SA. Mechanical state transitions in the regulation of tissue form and function. Nat Rev Mol Cell Biol 2024;25(8):654–70. 10.1038/s41580-024-00719-x.

Mescher M, Jeong P, Knapp SK, Rubsam M, Saynisch M, Kranen M, et al. The epidermal polarity protein Par3 is a non-cell autonomous suppressor of malignant melanoma. J Exp Med 2017;214(2):339–58. 10.1084/jem.20160596.

Mezzano V, Sheikh F. Cell-cell junction remodeling in the heart: possible role in cardiac conduction system function and arrhythmias? Life Sci 2012;90(9-10):313–21. 10.1016/j.lfs.2011.12.009.

Nanavati BN, Noordstra I, Lwin AKO, Brooks JW, Rae J, Parton RG, et al. The desmosome-intermediate filament system facilitates mechanotransduction at adherens junctions for epithelial homeostasis. Curr Biol 2024;34(17):4081–90 e5. 10.1016/j.cub.2024.07.074.

Nava MM, Miroshnikova YA, Biggs LC, Whitefield DB, Metge F, Boucas J, et al. Heterochromatin-Driven Nuclear Softening Protects the Genome against Mechanical Stress-Induced Damage. Cell 2020;181(4):800–17 e22. 10.1016/j.cell.2020.03.052.

Nekrasova O, Harmon RM, Broussard JA, Koetsier JL, Godsel LM, Fitz GN, et al. Desmosomal cadherin association with Tctex-1 and cortactin-Arp2/3 drives perijunctional actin polymerization to promote keratinocyte delamination. Nat Commun 2018;9(1):1053. 10.1038/s41467-018-03414-6.

Noethel B, Ramms L, Dreissen G, Hoffmann M, Springer R, Rubsam M, et al. Transition of responsive mechanosensitive elements from focal adhesions to adherens junctions on epithelial differentiation. Mol Biol Cell 2018;29(19):2317–25. 10.1091/mbc.E17-06-0387.

Paller AS, Teng J, Mazereeuw-Hautier J, Hernandez-Martin A, Granier Tournier C, Hovnanian A, et al. Syndromic epidermal differentiation disorders: a new classification toward pathogenesis-based therapy. Br J Dermatol 2025;193(4):592–618. 10.1093/bjd/ljaf123.

Parrish EP, Steart PV, Garrod DR, Weller RO. Antidesmosomal monoclonal antibody in the diagnosis of intracranial tumours. J Pathol 1987;153(3):265–73. 10.1002/path.1711530311.

Perl AL, Koetsier JL, Green KJ. PP2A-B55alpha controls keratinocyte adhesion through dephosphorylation of the Desmoplakin C-terminus. Sci Rep 2023;13(1):12720. 10.1038/s41598-023-37874-8.

Perl AL, Pokorny JL, Green KJ. Desmosomes at a glance. J Cell Sci 2024;137(12). 10.1242/jcs.261899.

Peskoller M, Bhosale A, Gobel K, Lohr J, Miceli S, Perot S, et al. How to Build and Regenerate a Functional Skin Barrier: The Adhesive and Cell Shaping Travels of a Keratinocyte. J Invest Dermatol 2022;142(4):1020–5. 10.1016/j.jid.2021.12.034.

Price AJ, Cost AL, Ungewiss H, Waschke J, Dunn AR, Grashoff C. Mechanical loading of desmosomes depends on the magnitude and orientation of external stress. Nat Commun 2018;9(1):5284. 10.1038/s41467-018-07523-0.

Protonotarios N, Tsatsopoulou A. Naxos disease and Carvajal syndrome: cardiocutaneous disorders that highlight the pathogenesis and broaden the spectrum of arrhythmogenic right ventricular cardiomyopathy. Cardiovasc Pathol 2004;13(4):185–94. 10.1016/j.carpath.2004.03.609.

Quinlan RA, Schwarz N, Windoffer R, Richardson C, Hawkins T, Broussard JA, et al. A rim-and-spoke hypothesis to explain the biomechanical roles for cytoplasmic intermediate filament networks. J Cell Sci 2017;130(20):3437–45. 10.1242/jcs.202168.

Rathod M, Franz H, Beyersdorfer V, Wanuske MT, Leal-Fischer K, Hanns P, et al. DPM1 modulates desmosomal adhesion and epidermal differentiation through SERPINB5. J Cell Biol 2024;223(4). 10.1083/jcb.202305006.

Rubsam M, Mertz AF, Kubo A, Marg S, Jungst C, Goranci-Buzhala G, et al. E-cadherin integrates mechanotransduction and EGFR signaling to control junctional tissue polarization and tight junction positioning. Nat Commun 2017;8(1):1250. 10.1038/s41467-017-01170-7.

Rubsam M, Pullen R, Tellkamp F, Bianco A, Peskoller M, Bloch W, et al. Polarity signaling balances epithelial contractility and mechanical resistance. Sci Rep 2023;13(1):7743. 10.1038/s41598-023-33485-5.

Sadhanasatish T, Augustin K, Windgasse L, Chrostek-Grashoff A, Rief M, Grashoff C. A molecular optomechanics approach reveals functional relevance of force transduction across talin and desmoplakin. Sci Adv 2023;9(25):eadg3347. 10.1126/sciadv.adg3347.

Schindelin J, Arganda-Carreras I, Frise E, Kaynig V, Longair M, Pietzsch T, et al. Fiji: an open-source platform for biological-image analysis. Nat Methods 2012;9(7):676–82. 10.1038/nmeth.2019.

Schmidt A, Koch PJ. Desmosomes: just cell adhesion or is there more? Cell Adh Migr 2007;1(1):28–32. 10.4161/cam.1.1.4204.

Seeley LD, Ainslie CM, Sewell-Loftin MK, Mattheyses AL. Super-resolution imaging reveals stretch-induced architectural rearrangement of desmoplakin in desmosomes. J Invest Dermatol 2026. 10.1016/j.jid.2026.04.016.

Soffer A, Bhosale A, Ghodrat R, Peskoller M, Matsui T, Niessen CM, et al. Spectrin coordinates cell shape and signaling essential for epidermal differentiation. J Cell Biol 2026;225(4). 10.1083/jcb.202502071.

Stappenbeck TS, Lamb JA, Corcoran CM, Green KJ. Phosphorylation of the desmoplakin COOH terminus negatively regulates its interaction with keratin intermediate filament networks. J Biol Chem 1994;269(47):29351–4.

Villeneuve C, Hashmi A, Ylivinkka I, Lawson-Keister E, Miroshnikova YA, Perez-Gonzalez C, et al. Mechanical forces across compartments coordinate cell shape and fate transitions to generate tissue architecture. Nat Cell Biol 2024;26(2):207–18. 10.1038/s41556-023-01332-4.

Yoshida K, Yokouchi M, Nagao K, Ishii K, Amagai M, Kubo A. Functional tight junction barrier localizes in the second layer of the stratum granulosum of human epidermis. J Dermatol Sci 2013;71(2):89–99. 10.1016/j.jdermsci.2013.04.021.

Zarkoob H, Kam CY, Koetsier JL, McCarthy E, Jaiganesh A, Kelsell DP, et al. RhoGEF Ect2 supports RhoA activity at cell-cell junctions through desmoplakin. Life Sci Alliance 2026;9(6). 10.26508/lsa.202503454.

