## Supplemental Figure 1 for "Regulation of the desmosome-intermediate filament linkage enables an adaptive mechano-response within the stratified epidermis"

A

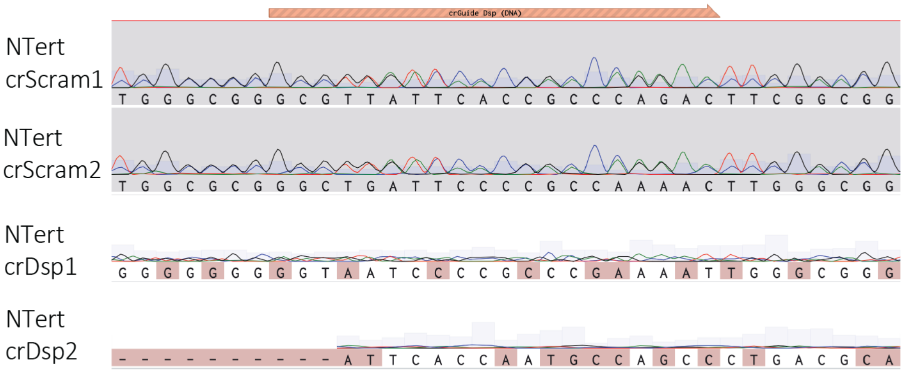


B

| | # of mismatches | Pairwise identity (%) |
|---|---|---|
| NTert crScram1 | 19 | 77 |
| NTert crScram2 | 11 | 79 |
| NTert crDsp1 | 102 | 66 |
| NTert crDsp2 | 196 | 28 |

C

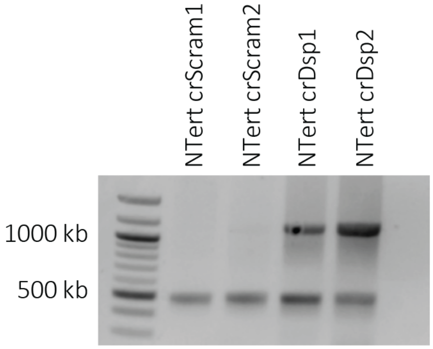


D

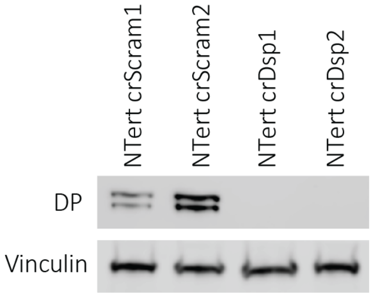
