## Supplementary figures and images for "Regulation of the desmosome-intermediate filament linkage enables an adaptive mechano-response within the stratified epidermis"

### Supplemental Figure 2

Supplemental Figure 2

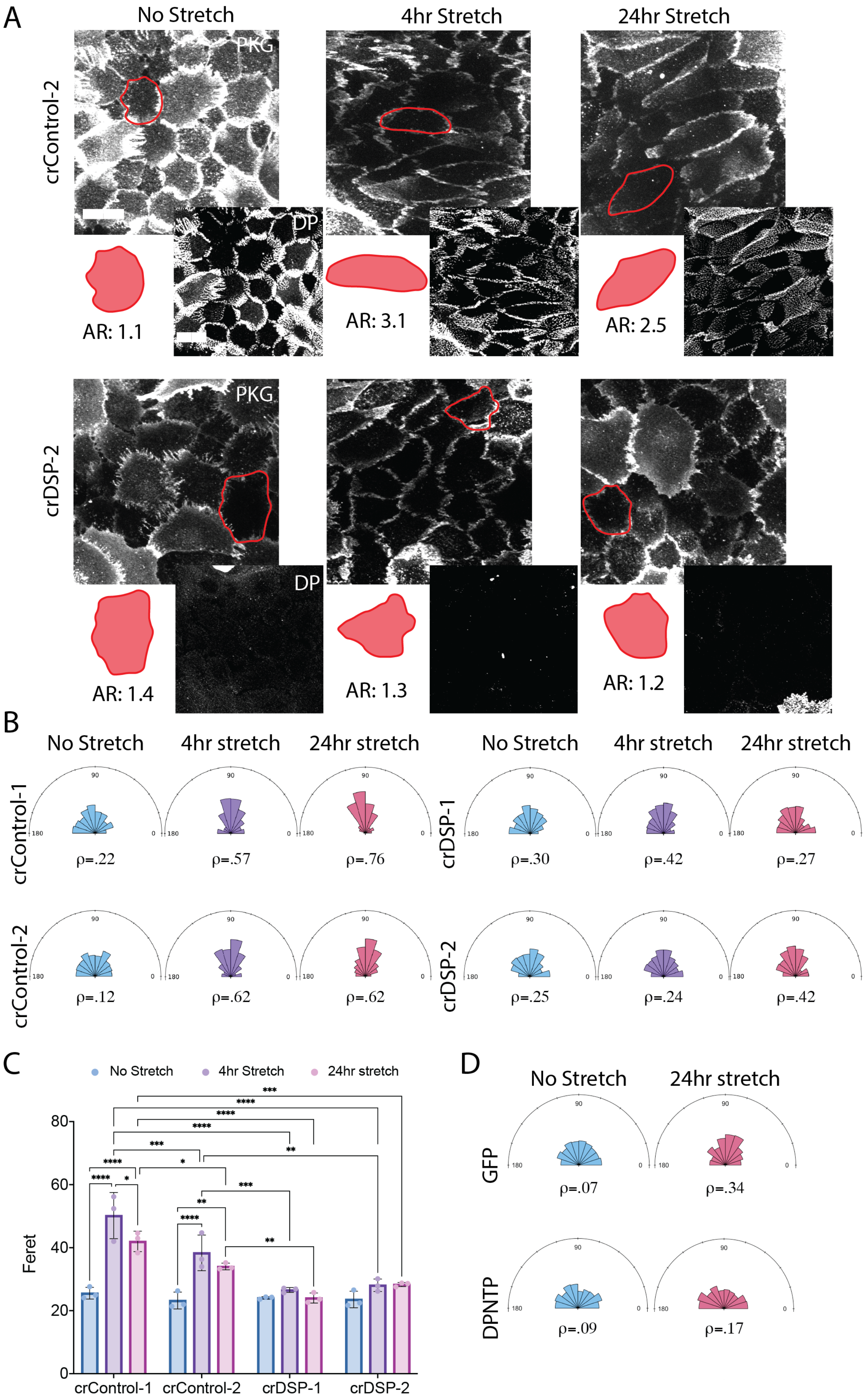

### Supplemental Figure 3

###### Supplemental Figure 3

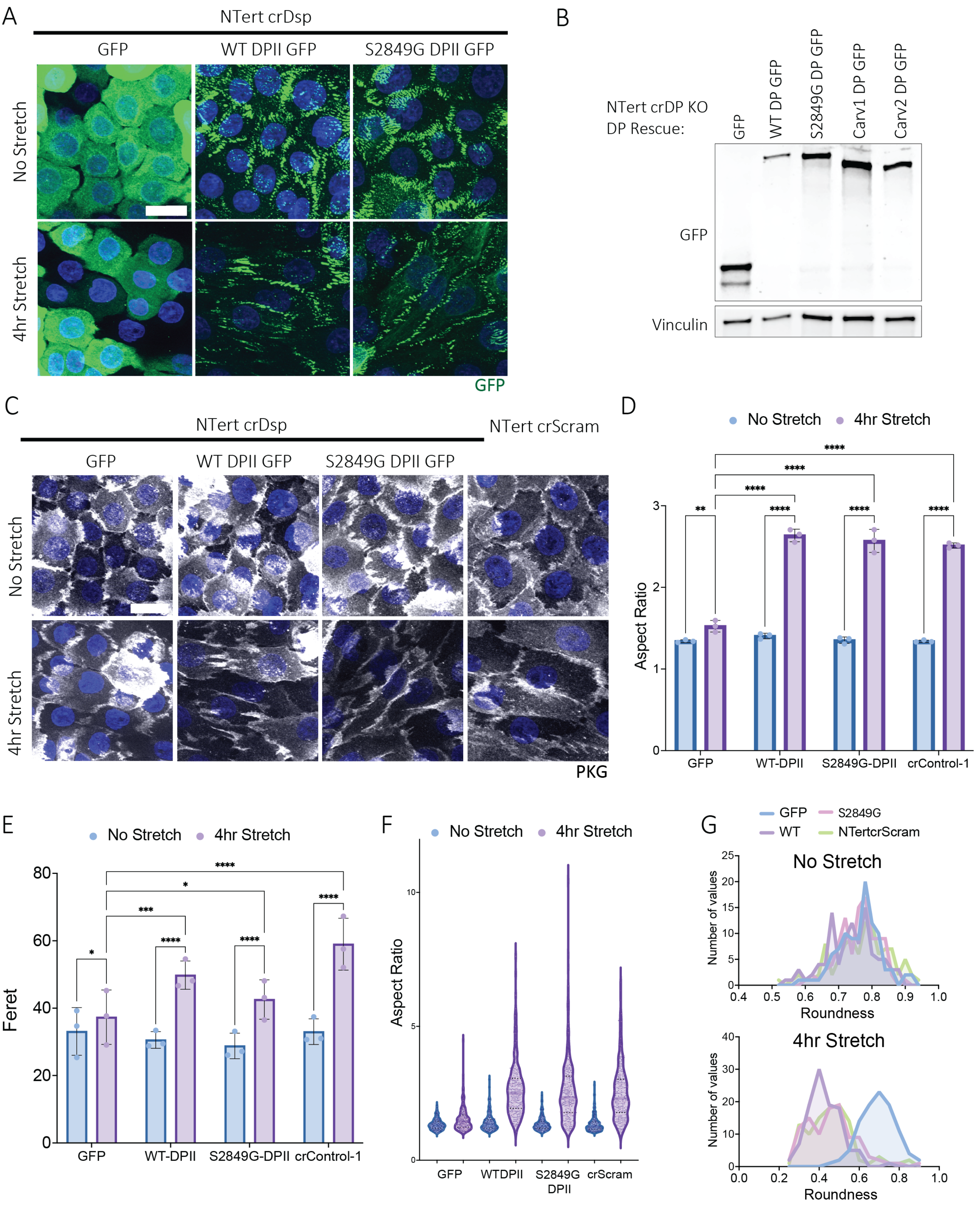

### Supplemental Figure 4

Supplemental Figure 4

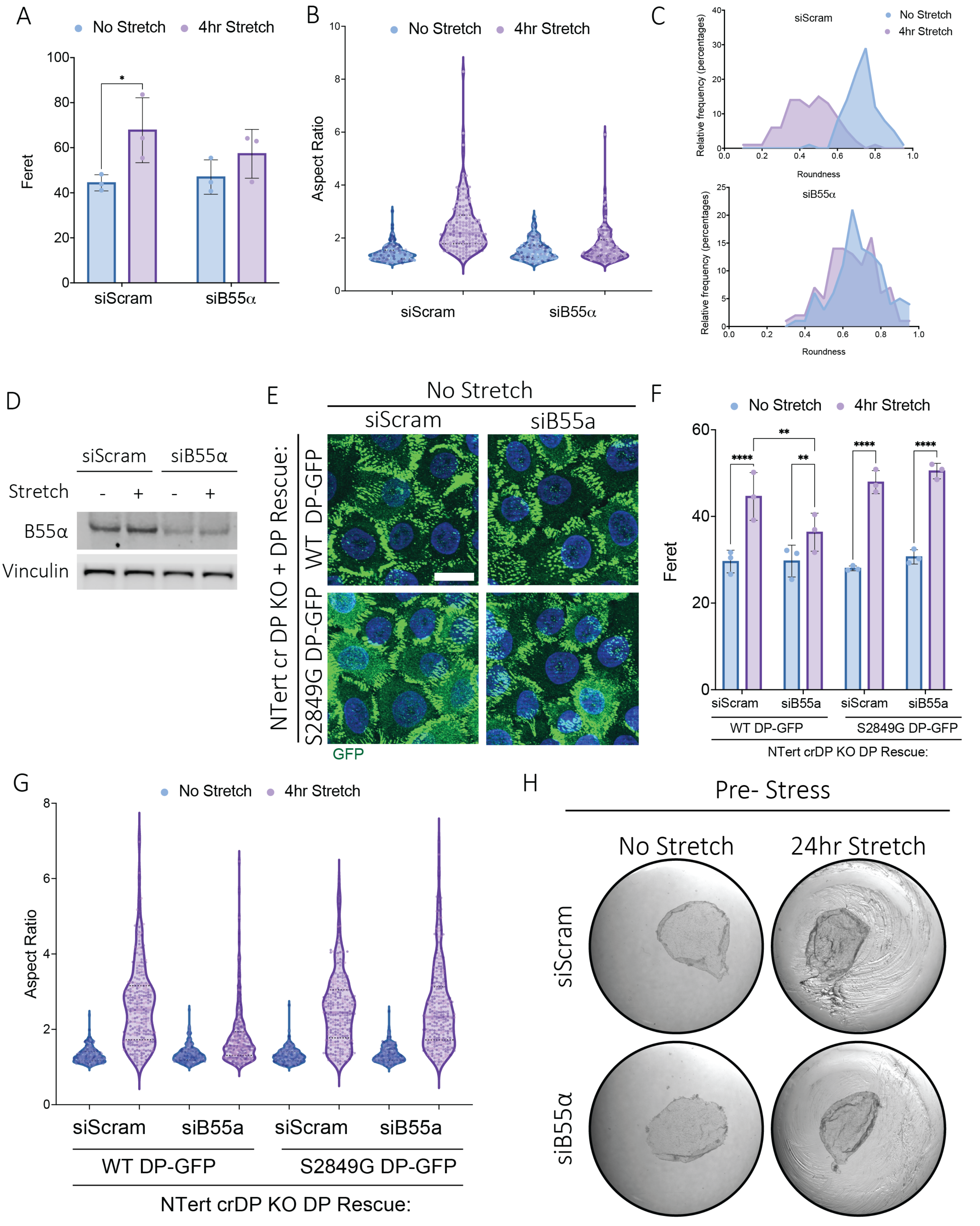

### Supplemental Figure 5

Supplemental Figure 5

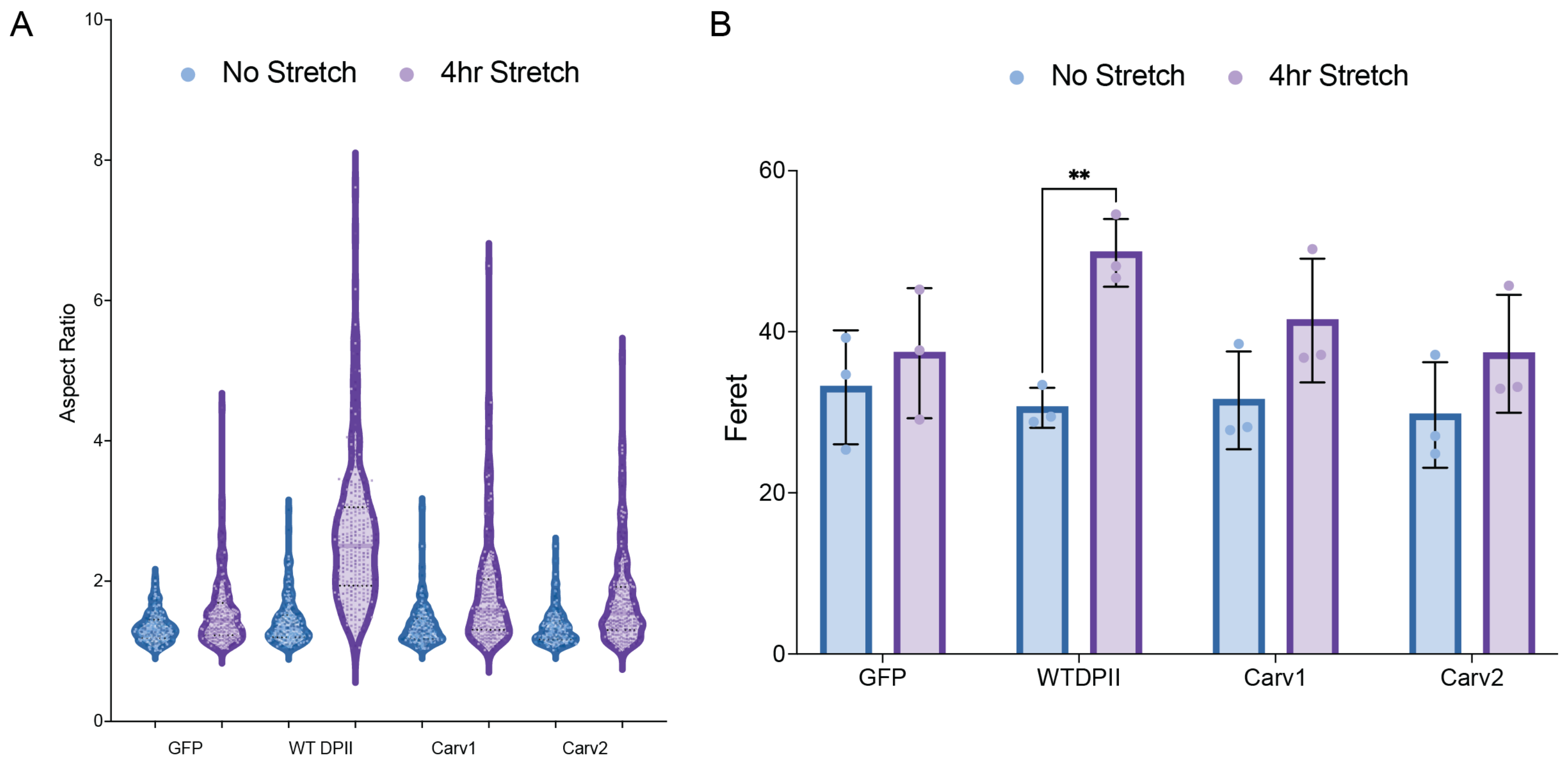

### Supplemental Movie 1

#### Slide 1
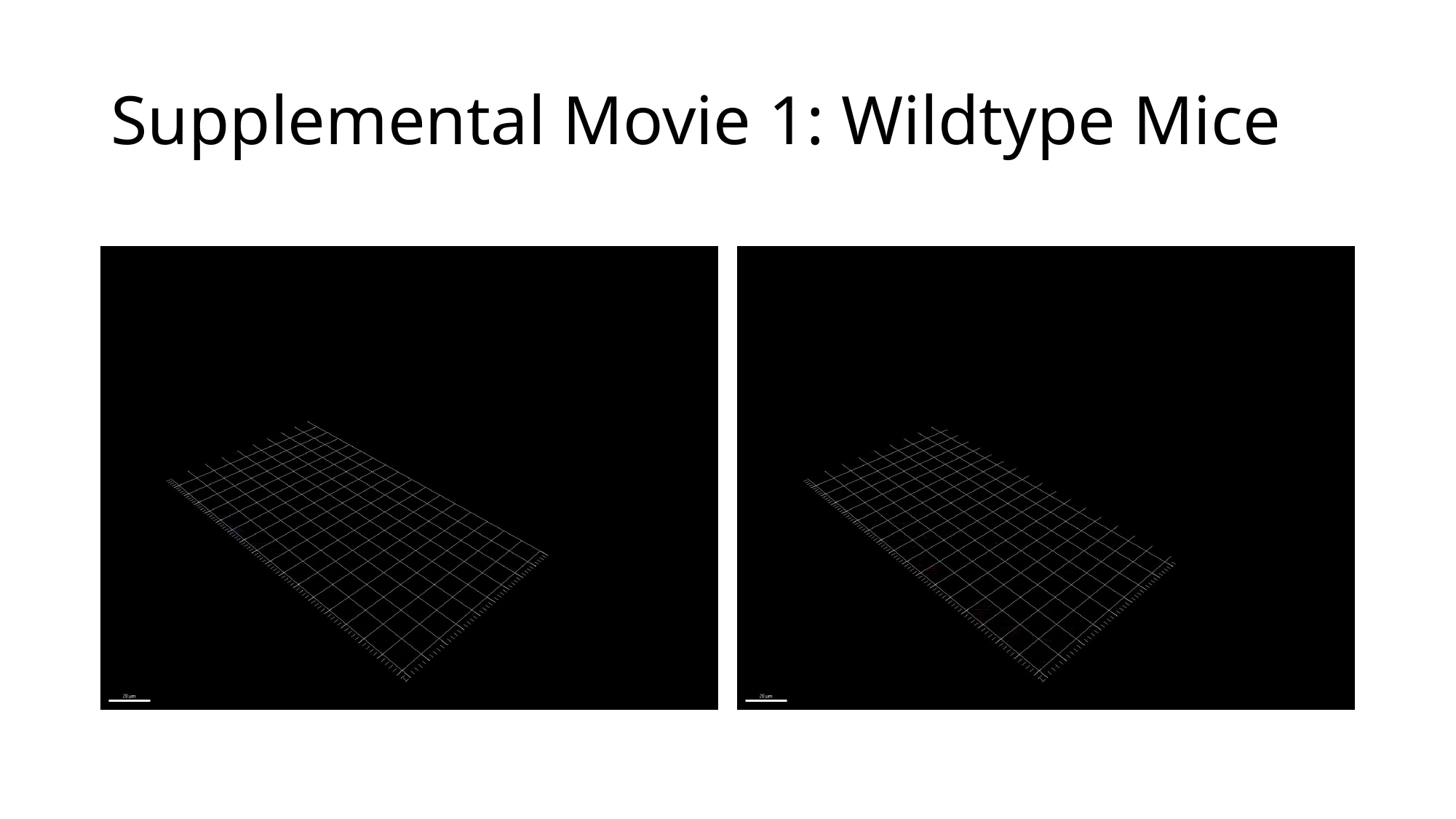

##### Supplemental Movie 1: Wildtype Mice

### Supplemental Movie 2

#### Slide 1
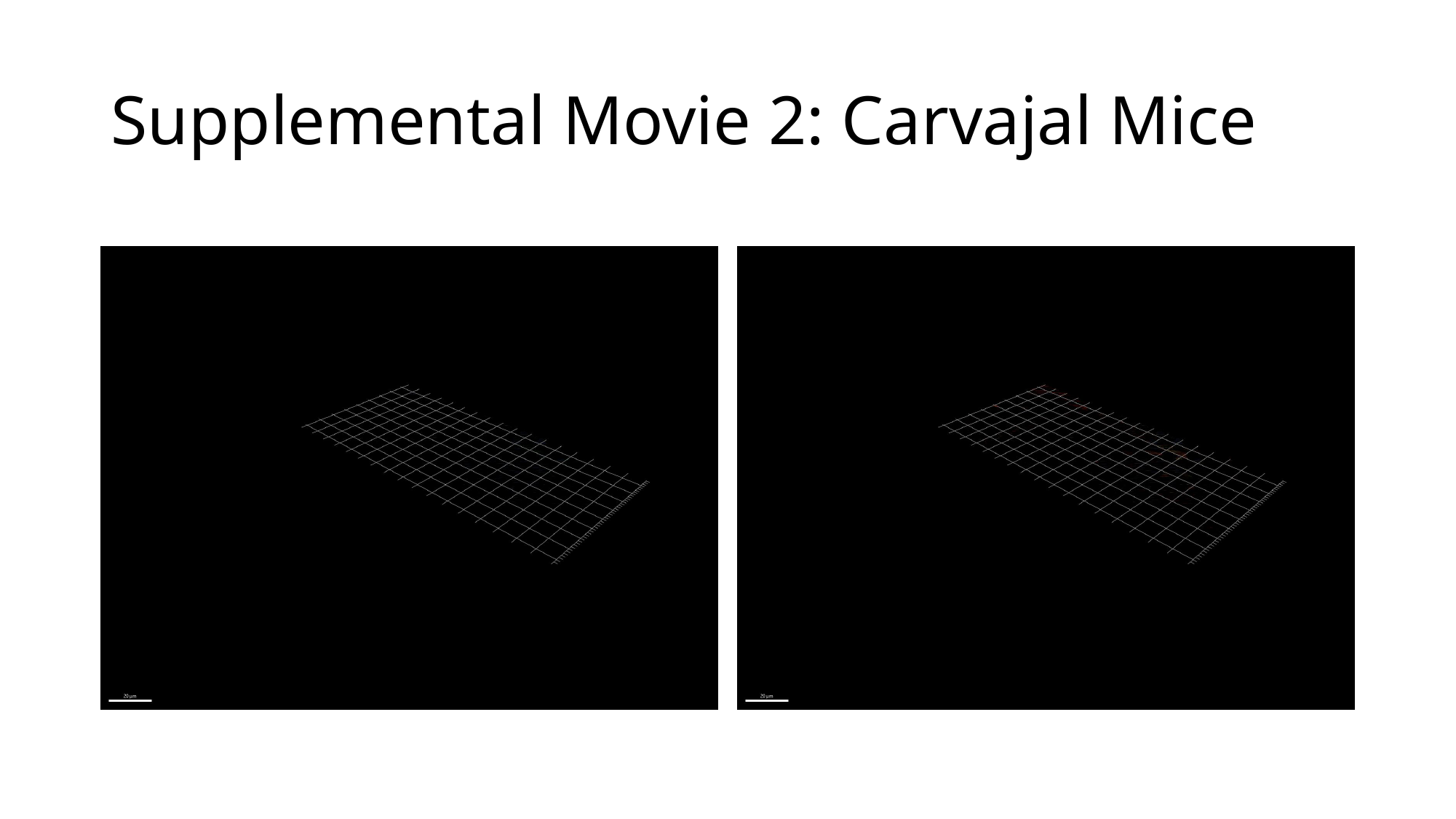

##### Supplemental Movie 2: Carvajal Mice
